# Network-based approach elucidates critical genes in BRCA subtypes and chemotherapy response in Triple Negative Breast Cancer

**DOI:** 10.1101/2023.05.21.541618

**Authors:** Piyush Agrawal, Navami Jain, Vishaka Gopalan, Annan Timon, Arashdeep Singh, Padma S Rajagopal, Sridhar Hannenhalli

**Author notes:** Corresponding authors Email: Sridhar Hannenhalli Piyush Agrawal. Equal Correspondence (Piyush Agrawal, Sridhar Hannenhalli).

## Abstract

Breast cancers exhibit substantial transcriptional heterogeneity, posing a significant challenge to the prediction of treatment response and prognostication of outcomes. Especially, translation of TNBC subtypes to the clinic remains a work in progress, in part because of a lack of clear transcriptional signatures distinguishing the subtypes. Our recent network-based approach, PathExt, demonstrates that global transcriptional changes in a disease context are likely mediated by a small number of key genes, and these mediators may better reflect functional or translationally relevant heterogeneity. We apply PathExt to 1059 BRCA tumors and 112 healthy control samples across 4 subtypes to identify frequent, key-mediator genes in each BRCA subtype. Compared to conventional differential expression analysis, PathExt-identified genes (1) exhibit greater concordance across tumors, revealing shared as well as BRCA subtype-specific biological processes, (2) better recapitulate BRCA-associated genes in multiple benchmarks, and (3) exhibit greater dependency scores in BRCA subtype-specific cancer cell lines. Single cell transcriptomes of BRCA subtype tumors reveal a subtype-specific distribution of PathExt-identified genes in multiple cell types from the tumor microenvironment. Application of PathExt to a TNBC chemotherapy response dataset identified TNBC subtype-specific key genes and biological processes associated with resistance. We described putative drugs that target top novel genes potentially mediating drug resistance. Overall, PathExt applied to breast cancer refines previous views of gene expression heterogeneity and identifies potential mediators of TNBC subtypes, including potential therapeutic targets.

## Introduction

Breast cancer (BRCA) is one of the leading cancers among women worldwide^1^. In 2023, 300,500 are expected to be diagnosed and 43,700 are expected to die from the disease [https://www.cancer.org/cancer/breast-cancer/about/how-common-is-breast-cancer.html]. Triple negative breast cancers (TNBC), defined clinically by the lack of estrogen receptor (ER), progesterone receptor (PR), or human epidermal growth factor receptor (HER2) on immunohistochemistry^2^ are especially aggressive with limited therapeutic options. Gene expression is used, primarily in the translational context, to characterize BRCA subtypes: Luminal A and Luminal B (which overlap with hormone-receptor positive breast cancers), HER2-enriched (which overlaps with HER2-positive tumors), and Basal-like (which overlaps with TNBC)^3,4^.

To identify potential targets for BRCA treatment, previous studies have relied on differentially expressed genes (DEGs). However, individual genes are subject to stochastic variability in gene expression, limiting their reproducibility and consistency^5^. Furthermore, transcriptional changes are often due to perturbations of key mediators (such as transcription factors, kinases, and other regulatory proteins) in a complex gene regulatory network. Identifying these potential mediators is more likely to provide mechanistic insights and therapeutic targets. Network-based models have been shown to improve potential target identification in various cancers^6–9^.

In contrast with DEGs, our previously developed network-based tool PathExt identifies differentially active pathways in a knowledge-based network and the corresponding central genes in the subnetwork (called TopNet) composed of the differentially active paths. Central genes identified by PathExt offer a more robust and mechanistic view of transcriptional changes across conditions compared to DEGs, explaining differential expression of downstream genes^10^.

We applied PathExt framework to the TCGA BRCA transcriptional data (tumor and normal samples)^11^ to identify key genes mediating the global transcriptional changes in each BRCA subtype. Against multiple benchmark datasets, PathExt-identified genes substantially outperform DEGs in recapitulating potential driver genes and performs favorably compared to another network-based approach, MOMA^12^. PathExt-identified genes are further validated by their effect on cellular viability, based on CRISPR knock-out data from the DepMap database^13,14^, as well as their greater-than-expected mutation frequencies in TCGA samples, in a subtype-specific manner (note that the PathExt relies only on the transcriptome data and does not utilize the mutational data). We further applied PathExt in a clinical trial of neoadjuvant chemotherapy in TNBC to identify key genes associated with treatment-resistant tumors. Lastly, based on computational drug screening, we propose potential therapeutic strategies for targets identified by PathExt. Overall, PathExt refined prior characterization of inter-tumor heterogeneity and identified more consistent genes associated with BRCA subtypes, as well as candidate genes to study resistance to neoadjuvant chemotherapy in TNBC.

## Results

### Overview of the workflow

The overall workflow of the study is shown in **Figure 1**. Given a BRCA transcriptomic profile and control samples, as well as a knowledge-based protein interaction network, PathExt^10^ identifies key genes likely to mediate the observed global transcriptomic changes in the specific sample relative to the control samples. We separately identify genes mediating up-regulation (based on Activated TopNet) as well as down-regulation (based on Repressed TopNet) of gene expression. We applied PathExt to 1059 TCGA BRCA transcriptomic samples across 4 BRCA subtypes using 112 healthy controls to identify top 100 key mediator genes in each sample (Methods). We further identified the top 200 genes most frequently identified across samples in each cancer subtype as a key mediator. As a control, we followed an analogous procedure to identify the top 200 most frequent up and downregulated DEGs in each subtype. Complete lists of genes and their frequencies are provided in the **Supplementary Table S1-S4**. We performed a series of downstream analyses to assess the relative merits of these identified genes in terms of their functional roles in BRCA. We additionally analyzed a TNBC dataset to identify central genes associated with nonresponse to neoadjuvant chemotherapy in various TNBC subtypes.

**Figure 1:**
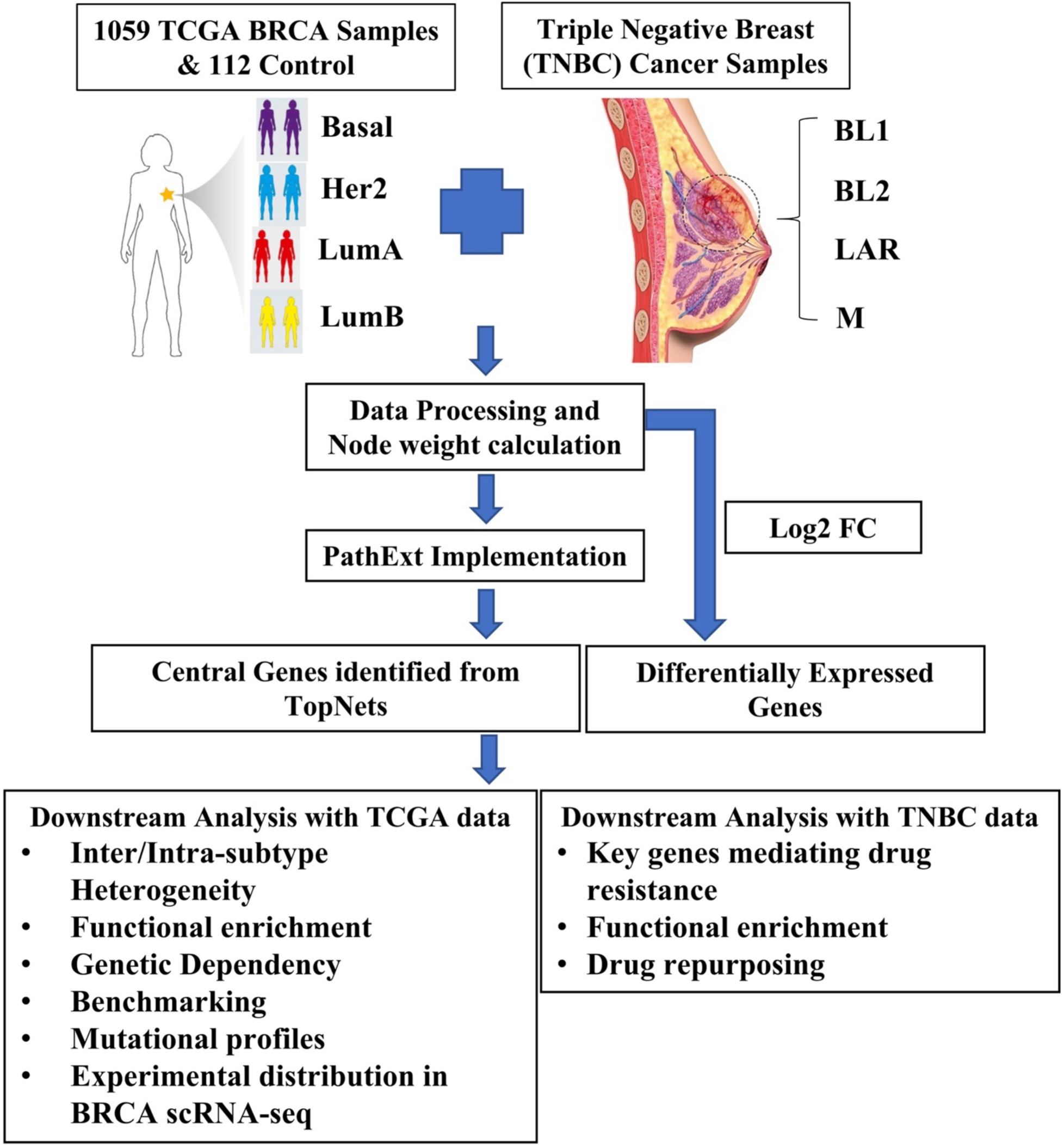
Architecture of the study. PathExt was implemented on the TCGA-BRCA dataset to identify subtype-specific most frequent central mediating genes as well as DEGs for comparison. Central genes were used for various analyses such as identifying enriched biological processes, genetic dependencies of the cell, identifying cell types mediating global transcriptomic response in single cell data, recapitulating breast cancer driver genes, FDA approved breast cancer targets, comparison with previous network-based methods and DEGs. PathExt was also implemented on the TNBC dataset to identify key genes associated with non-response to neoadjuvant chemotherapy treatment in subtype-specific manner.

### PathExt reveals putative BRCA subtype-specific and shared mechanisms more consistently than DEGs

The following analyses are based on the PathExt-identified top 200 most frequent central genes in Activated and Repressed TopNets for each subtype (**Supplementary Table S5)** and the analogously calculated top 200 most frequent upregulated and downregulated DEGs (**Supplementary Table S6)**.

PathExt genes are much more frequently represented within subtype-specific samples than DEGs, suggesting that PathExt may better reveal shared mechanisms across tumors. For example, in the Basal subtype, top Activated PathExt genes *CDC20* and *TTK* were key genes in 175/177(∼99%) tumors, while the most frequently upregulated DEG - *CT83* was the top DEG in only 117/177(∼66%) tumors. Likewise, in the HER2 subtype, out of 80 samples, PathExt identified *ERBB2* (encoding HER2) as the central gene in 42 samples (>50%) whereas it was differentially upregulated in only 11 patients (∼15%), underscoring the potential of PathExt in capturing functionally relevant genes.

PathExt genes are also much more frequently represented than DEGs when comparing across all breast cancer samples, as shown in **Figure 2A** for Activated TopNets and **Figure 2B** for Repressed TopNets. Direct comparison of genes prioritized by PathExt and DEGs also revealed substantial differences **(Supplementary Figure S1 A&B)**.

**Figure 2:**
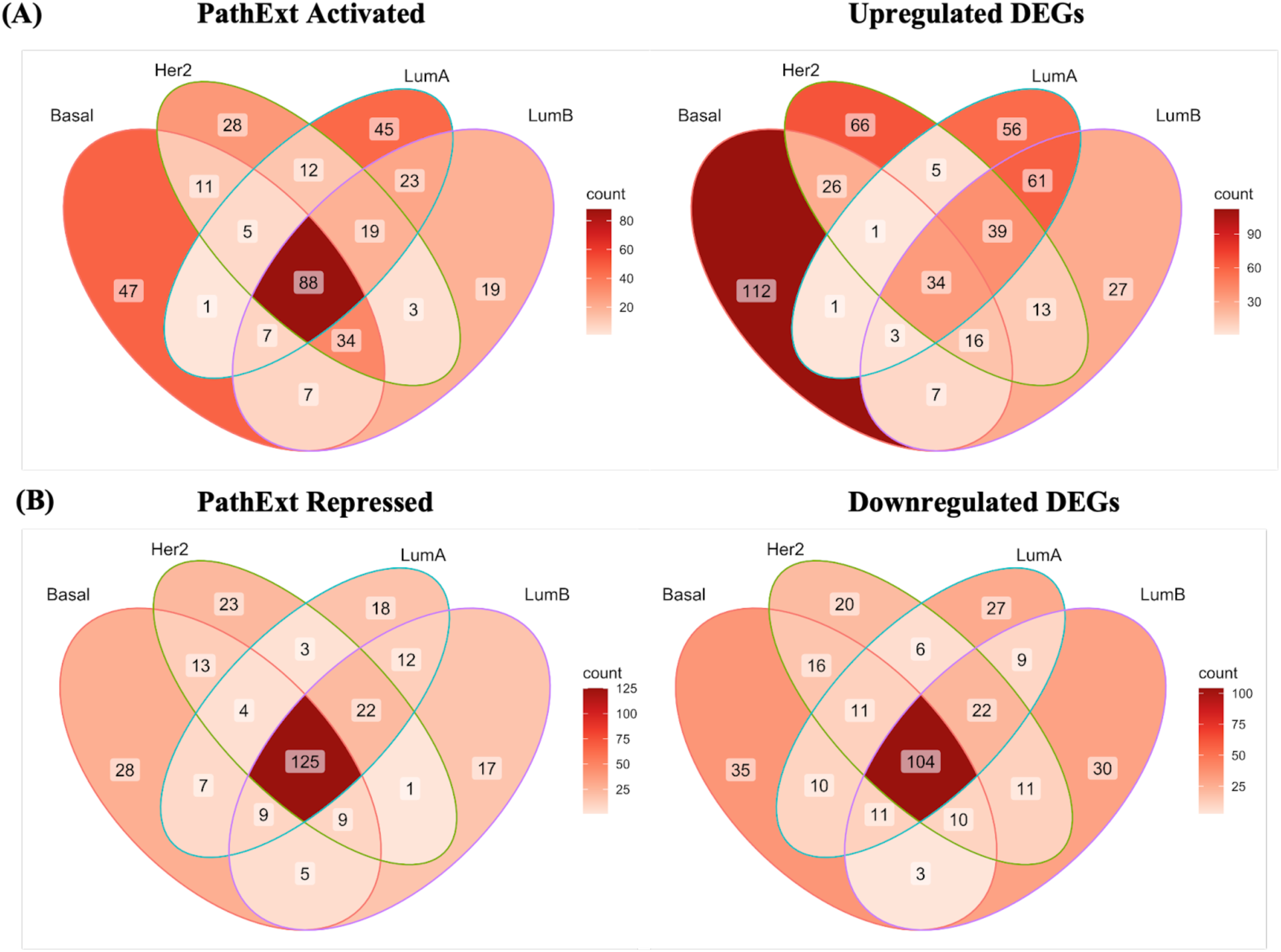
Venn diagram showing gene overlap among central genes in various subtypes identified in (A) PathExt Activated TopNets and Upregulated DEGs and for (B) PathExt Repressed TopNets and Downregulated DEGs.

We next identified the enriched biological processes associated with pan-subtype and subtype-specific genes. For the PathExt Activated TopNets, as expected, the common genes were associated mostly with cell cycle **(Figure 3A, Supplementary Table S7)**. However, subtype-specific genes were enriched for distinct processes that are largely supported by literature. For instance, the Basal subtype was associated with cell fate determination, regulation of T-cell proliferation, neuron differentiation, interferon-gamma production, etc.^15–18^ **[Figure 3B]**; enriched processes for other subtypes are summarized in **Supplementary Figure S2A-B** (except LumB, where no enrichment was found), and the complete list of enriched processes is provided in the **Supplementary Table S8-S10**. Among top genes in Repressed TopNets, most subtypes share similar processes, including neuropeptide signaling pathway, cellular hormone metabolic pathway, muscle cell development, etc. **(Figure 3C)** whereas unique genes for the Basal subtype were enriched for the processes such as terpenoid metabolic process, liver development, etc. **(Figure 3D)**. For remaining subtypes see **Supplementary Figure S2C-E** and the complete list of enriched processes are provided in the **Supplementary Table S11-S15**. In contrast, when analyzing pan-subtype and subtype-specific DEGs, distinct, but fewer, sets of processes were enriched. Notably, for many subtypes we didn’t observe any significant enriched processes associated with up and downregulated DEGs; all results are provided in **Supplementary Figure S3, Supplementary Table S16-S21.**

**Figure 3:**
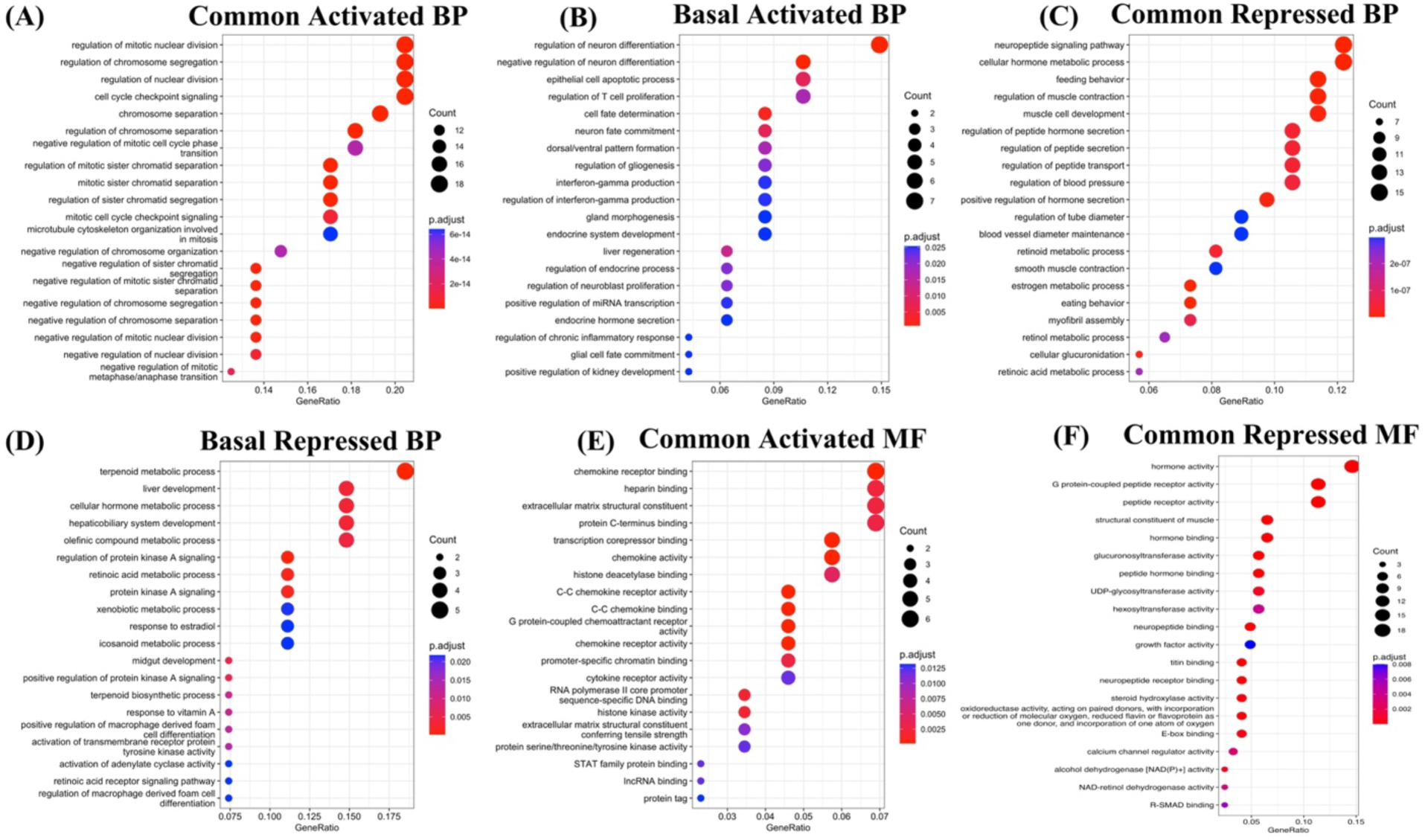
Top central PathExt central genes are associated with essential biological processes and molecular functions. Top20 enriched biological processes associated with activated common central genes among all the 4 BRCA subtypes (A); Top20 enriched biological processes associated with activated genes uniquely in Basal (B); Top20 enriched biological processes associated with repressed common central genes among all the 4 BRCA subtypes (C); and Top20 enriched biological processes associated with repressed genes uniquely in Basal (D); Top20 enriched molecular functions associated with activated common central genes among all the 4 BRCA subtypes (E); and, Top20 enriched molecular functions associated with repressed common central genes among all the 4 BRCA subtypes (F).

Enriched molecular functions for Activated TopNets of pan-subtype genes include chemokine binding, extracellular matrix structural constituent, and transcription corepressor binding **(Figure 3E)**. Subtype-specific genes (except Her2, where no enrichment was found) also revealed subtype-specific molecular functions supported by literature^19–22^ **(Supplementary Figure S4A-C, Supplementary Table S22-S25).** Likewise for the Repressed TopNets, molecular functions enriched among pan-subtype genes include hormone activity, glycosyltransferase activity, and titin binding. **(Figure 3F).** Enriched functions among subtype-specific PathExt genes (except LumB, where no enrichment was found) included somewhat related functions, e.g., G protein-coupled peptide receptor activity^23^ **(Supplementary Figure S4D-F, Supplementary Table S26-S29).** Pan-subtype and subtype-specific DEGs revealed processes only partly overlapping with PathExt (**Supplementary Figure S5, Supplementary Table S30-S34).** Notably, for many subtypes we didn’t observe any significant enriched molecular functions associated with up and downregulated DEGs.

Overall, PathExt reveals greater commonality across samples and across subtypes, and identifies many shared and subtype-specific genes and processes mediating the global transcriptome shifts that are missed by the conventional DEG analysis, underscoring the complementary value of PathExt approach.

Next, we assessed the extent to which the subtype-specific genes identified by PathExt have any support for their subtype-specific functionality. We did this with respect to subtype-specific expression and mutational patterns of those genes. First, for every gene, in a given subtype, we computed the log-fold difference of that gene’s expression in the subtype relative to other subtypes. Unsurprisingly, subtype-specific central Activated genes show higher relative gene expression in the particular BRCA subtype, while central Repressed genes exhibit lower relative expression in the BRCA subtype **(Supplementary Figure S6)**. To characterize subtype-specific mutational patterns (which was not used by PathExt), for every gene, we separately computed the frequency of activating (copy number amplification; CNA) and inactivating (copy number loss, nonsense, frameshift indel) mutations in a subtype-specific manner. The mutation frequency was normalized by average frequency across all genes in a given BRCA subtype and further normalized by the same quantity for the other three subtypes to get a subtype-specific relative log-normalized mutation frequency. While the signals were weak owing to the sparsity of mutations, we broadly observed a subtype-specific enrichment of activating mutations in subtype-specific genes (both from Activated and Repressed TopNets) **(Figure 4A)** and a depletion of inactivating mutations **(Figure 4B).** Supported by previously reported dominance of CNAs in BRCA genomic landscape^24^, our results link CNAs to key genes affecting global transcriptome in a BRCA subtype-specific fashion. Overall, our analyses support the role of key genes identified by PathExt in BRCA subtype-specific pathogenesis.

**Figure 4:**
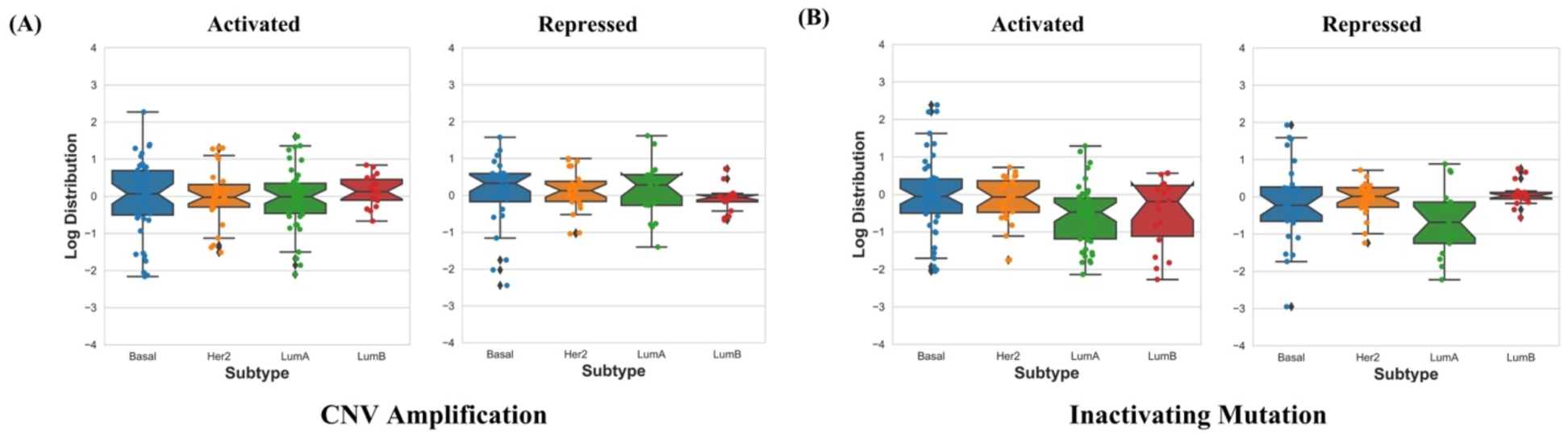
PathExt reveals subtype specific mutational properties. Boxplot representation of subtype-specific PathExt Activated and Repressed TopNets unique gene CNV amplification distribution (A); and subtype-specific PathExt Activated and Repressed TopNets unique gene inactivating mutation distribution (B).

### PathExt-identified genes perform better than DEGs in cell line, functional, and benchmarking tests

The key feature of PathExt is identification of genes that may not be overtly differentially expressed but may play a central role in mediating the tumor phenotype. First, we assess the extent to which subtype-specific PathExt genes exhibit subtype-specific function and clinical relevance. To do this, we utilized the publicly available DepMap dataset reporting cellular viability upon genome-wide CRISPR-Cas9 Knock Out (KO) across hundreds of cell lines, providing a dependency probability score for each gene in each cell line, where a higher score indicates greater essentiality or dependency^25^; by default, a score >0.5 indicates dependency. Critically, one can obtain dependency for each gene in BRCA subtype-specific cell lines. For each gene-subtype pair, we computed the gene’s average dependency probability scores across the cell lines derived from the specific BRCA subtype. We then compared the score distribution for PathExt genes with that for DEGs. As shown in **Figure 5A**, PathExt Activated genes exhibited significantly greater dependency in subtype-specific cell lines compared to upregulated DEGs. Interestingly, gene dependency of the Repressed TopNets and downregulated DEGs were very low suggesting that these genes do not affect cell survival. Our conclusions drawn from **Figure 5A** do not change when we compare the fraction of essential genes (dependency probability score > 0.5) between PathExt and DEGs based on Fisher’s test **(Supplementary Figure S7).** Depmap gene dependency probability value for the central genes (when available) of PathExt and DEGs in various subtypes is provided in the **Supplementary Table S35-38.**

**Figure 5:**
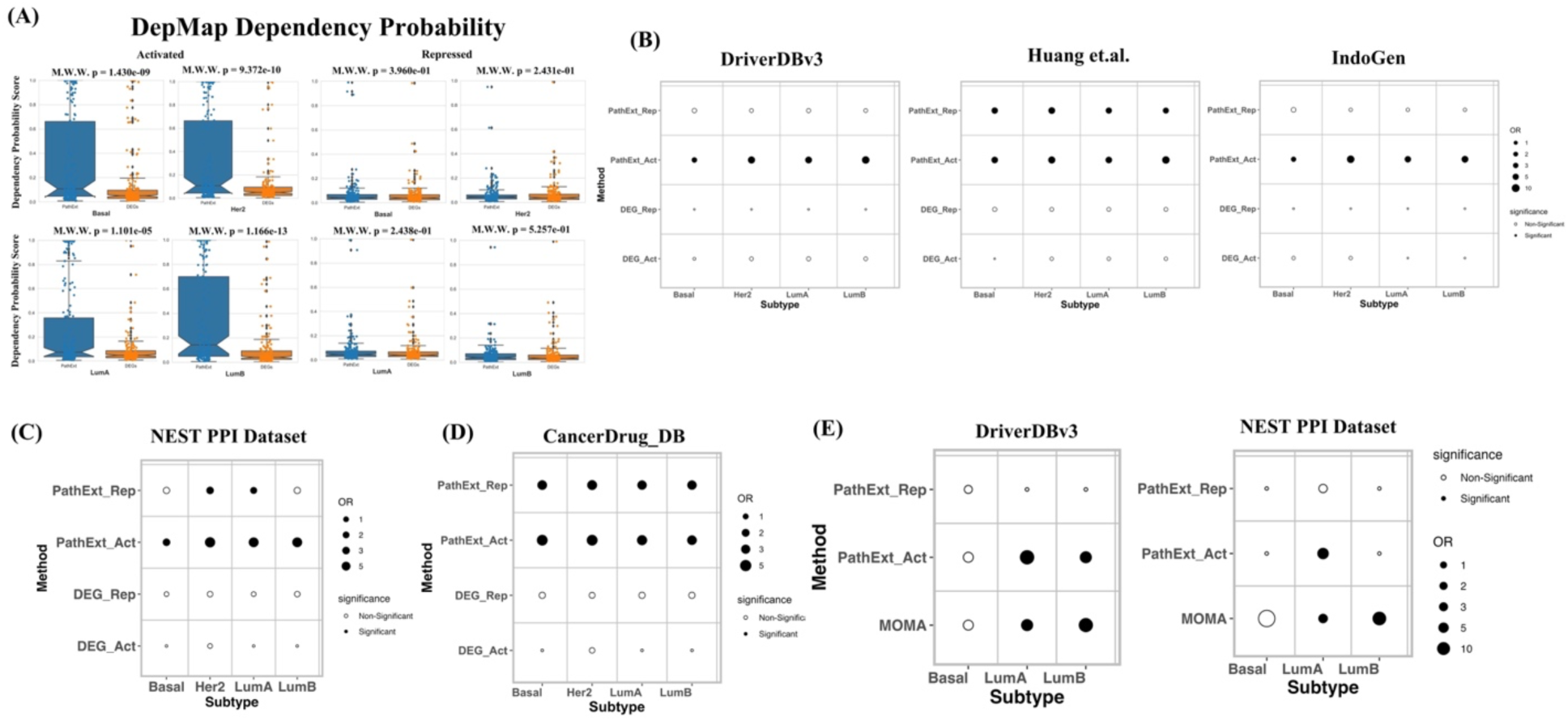
Evaluation of PathExt approach with DEGs and MOMA. Boxplot representation of gene dependency probability distribution by PathExt Activated TopNet genes and Upregulated DEGs and PathExt Repressed TopNet genes and Downregulated DEGs in various BRCA subtypes (A); Fishers Odds Ratio (OR) showing that PathExt recapitulated BRCA specific driver genes from multiple datasets (B); NEST PPI mutated modules comprising a number of genes affecting various mechanisms (C); targets associated with FDA approved BRCA drugs compared to DEGs with high statistical significance (D). PathExt showed comparable performance with MOMA (another network-based method) on various datasets (E).

Next, we assessed how significantly PathExt and DEGs recapitulated previously compiled driver genes in three 3 publicly available datasets: (i) DriverDBv3^26^ (including 313 BRCA driver genes) (ii) IntoGen^27^ (including 99 BRCA driver genes) and (iii) Huang et.al. dataset^28^ (including 500 BRCA driver genes). For each benchmark gene set, we assessed the significance of overlap (using Fisher Exact test) with the top 200 PathExt genes or DEGs in Activated or Repressed scenarios. As shown in **Figure 5B**, in all benchmarks, PathExt Activated TopNet central genes show significant overlap with the BRCA drivers, while upregulated DEGs do not.

Integrating protein interaction networks and tumor mutation profiles, Zheng et al. have reported a map of 395 protein systems (NEST) which are recurrently mutated in one or more cancer types likely under somatic mutation selection^29^. PathExt genes are associated significantly with several NEST systems, including Cell cycle, Nucleosome and Ribosome, Cytoplasm and Extracellular space, and Signaling Systems. More specifically, PathExt Activated TopNet central genes from all four subtypes showed significant overlap with NEST genes while central genes from Repressed TopNets showed significant overlap for subtypes Her2 and LumA. In sharp contrast, none of the DEGs (up or downregulated) from any of the subtypes showed significant overlap with the NEST systems **(Figure 5C)**.

Next, we compiled an additional translational benchmark gene set comprising 474 unique target genes of 46 FDA-approved BRCA drugs from CancerDrugs_DB^30^. Again, we found that these genes exhibit significantly higher overlap with the top 200 Activated PathExt genes compared to DEGs **(Figure 5D)**.

Finally, we compared PathExt with MOMA (Multi-omics Master-Regulator Analysis)^12^, another network-based approach which integrated transcription and mutational data in the context of inferred regulatory networks, and previously reported 407 master regulator (MR) proteins across 20 TCGA cohorts. These MR proteins are further grouped into 24 pan-cancer master regulator blocks. Specifically, the study has reported 39, 92 and 23 MR proteins for Basal, LumA and LumB subtypes, respectively. We compared the overlap of MOMA MR proteins with the DriverDBv3 driver cancer gene dataset and NEST PPI dataset with that of PathExt identified TopNet genes. For a fair comparison, we selected the same number of genes (based on frequency) which MOMA had for each subtype. Based on the Fisher test, PathExt genes exhibit a comparable overlap with the benchmark sets relative to MOMA **(Figure 5E)**.

Overall, PathExt identified genes shows greater cell dependency, recapitulates previously identified cancer drivers, BRCA specific mutated modules, and BRCA FDA approved drug targets far more effectively than the conventional DEG approach and performs favorably to a recent network based multi-omics approach MOMA.

### PathExt leverages single-cell data to identify cell-specific mediators in tumor micro-environment

Breast cancer heterogeneity is attributable to both transcriptional variation in cancer cells as well as variation in cellular composition in the tumor microenvironment. We therefore analyzed BRCA subtype-specific single cell transcriptomic data by Qian et. al.^31^ to chart the distribution of expression of the top 200 PathExt-identified genes across different cell types in the tumor microenvironment (Methods). Within each subtype-specific dataset we first identified in each cell type the genes that are expressed in that cell type with an above-mean expression; note that a gene can be expressed in multiple cell types, but not all. We then examined, in a subtype-specific manner, the fractions of PathExt genes expressed in each cell type relative to background expectation across all genes.

In the Qian et. al. dataset, the top genes from the Activated TopNets are expressed most frequently in the malignant cells, except in LumA, where top genes are most enriched in myeloid cells and depleted in malignant cells **(Figure 6A).** Additional cell types show enriched expression for top activated genes as well -- T cells in TNBC, myeloid cells in Her2, and B cells in LumB. Repressed central genes show similar broad trends except that in three subtypes central repressed genes are also enriched in fibroblasts, consistent with the enrichment of extracellular matrix processes among these genes **(Figure 6B)**.

**Figure 6:**
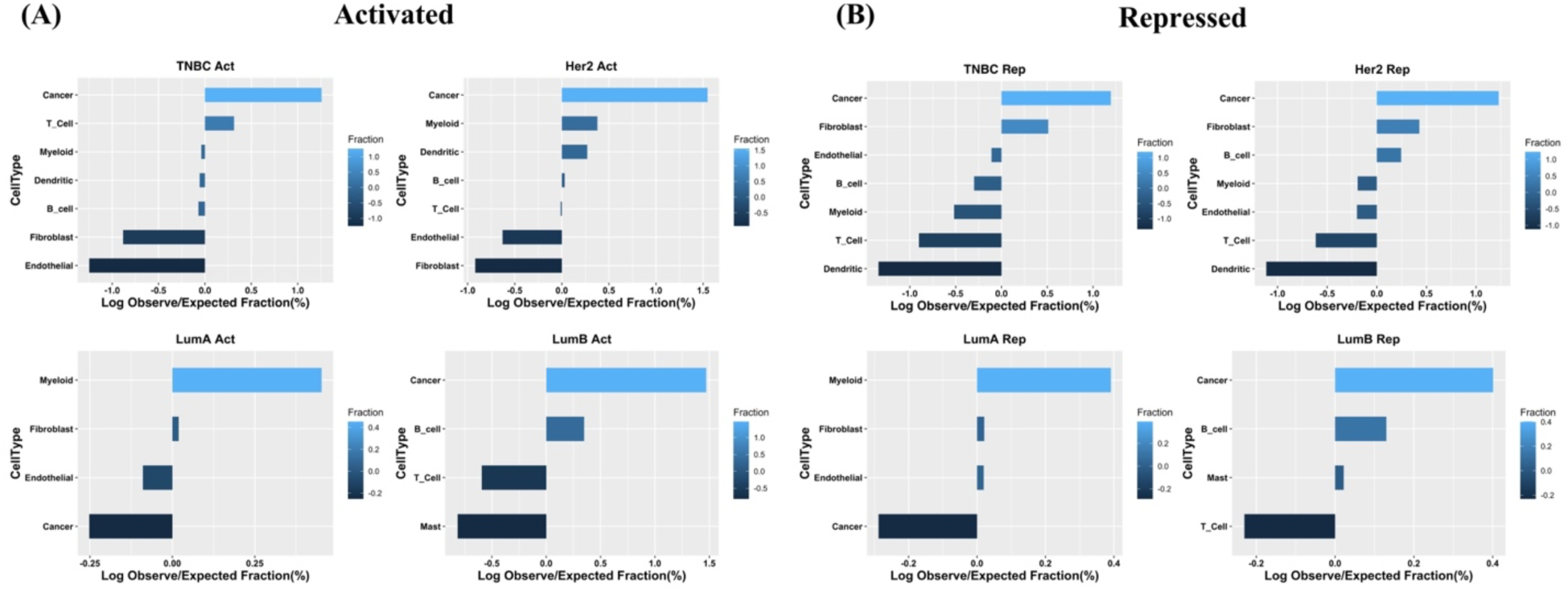
Gene expression distribution of top200 PathExt (A) Activated central genes and (B) Repressed central genes in individual cell types in each BRCA subtype. Within each subtype-specific scRNA-seq dataset, we first identified in each cell type the genes that are expressed in that cell type with an above-mean expression. We then examined the fractions of PathExt genes expressed in each cell type (Observed Frequency) relative to background expectation across all genes (Expected Frequency). Finally, Log (Observed/Expected) value was computed for the central genes across cell types.

### PathExt identifies potential mediators of resistance to Neoadjuvant Doxorubicin/Cyclophosphamide followed by Ixabepilone/Paclitaxel in TNBC

TNBC is the most aggressive BRCA subtype and presents a major therapeutic challenge. Although TNBC is a clinically defined subtype, it substantially overlaps with the Basal subtype defined based on transcriptional profile. Gene expression has been used to further classify TNBC breast cancers^32^. Here, we aim to apply PathExt to identify key genes potentially mediating therapeutic resistance. We used publicly available data from a phase II trial of neoadjuvant doxorubicin/cyclophosphamide followed by ixabepilone/paclitaxel therapy, where pathologic complete response (PCR) was the measure of response^33^. We specifically focused on the 138 TNBC samples classified into 4 subtypes (BL1-47, BL2-27, LAR-21, and M-43), further classified as responders or non-responders. We applied PathExt to identify central genes (Activated & Repressed) associated with non-responders (in each sample independently), using samples from responders as the control, and ranked genes based on number of samples of a specific TNBC subtype in which they were detected among the top 100 central genes. Complete lists of PathExt identified TNBC subtype-specific genes and their frequencies in Activated and Repressed TopNets associated with non-responders are provided in the **Supplementary Table S39 & S40 respectively.**

First, we selected the top 20 most frequent central genes in Activated TopNets from each TNBC subtype yielding 60 non-redundant genes, classified in 5 parts: Common - the genes present in at least 2 of the 4 TNBC subtypes among the top 20 genes, and 4 gene sets comprising genes uniquely found in a subtype among the top 20. **Figure 7** shows the subtype-specific frequencies of these genes; note that a gene, say *TGFB1*, may be among the top 20 uniquely in BL1 and yet may have a high frequency (although not among top 20) in another subtype (M). As can be seen, most subtypes show relatively high specificity among the top genes. **Supplementary Figure S8** shows a similar figure for top 20 central genes in Repressed TopNets in each subtype.

**Figure 7:**
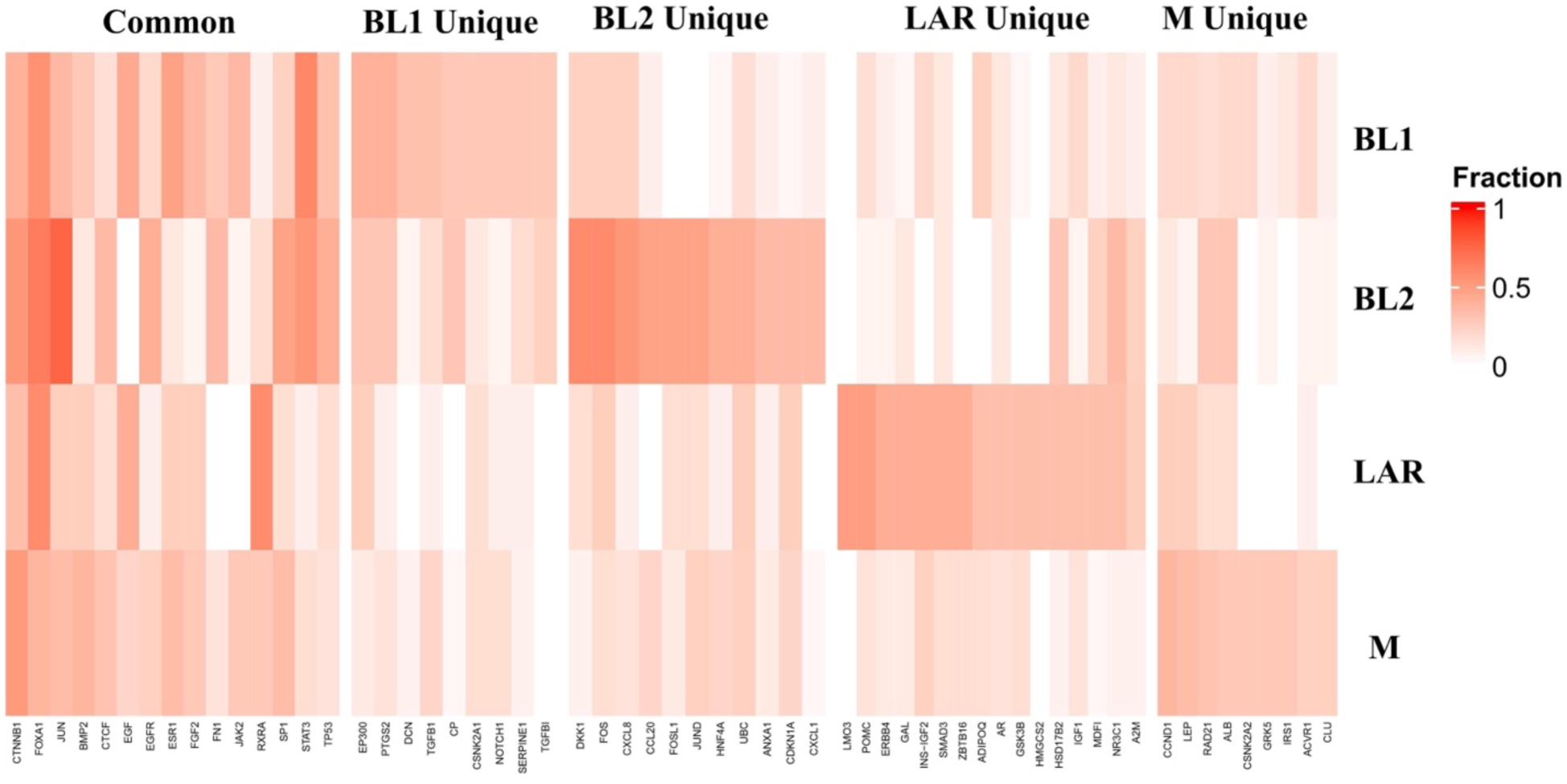
Top PathExt Activated unique and common gene fraction distribution in various TNBC subtypes. Top20 most frequent genes were selected from each TNBC subtype, from which pan-subtype common genes (top20 in at least two subtypes) and subtype-specific unique genes were identified. The heat plot shows, in each subtype, the fraction of samples in which the gene was among the top 20 central genes.

Next, we examined the top 100 most frequent central genes from each subtype and identified genes common among all subtypes and unique to a given subtype. For the activated TopNets, 5 genes -- *FOXA1, CTNNB1, JUN, FOS* and *ALB* were found in all TNBC subtypes and 49, 55, 61 and 48 genes were unique to BL1, BL2, LAR and M subtypes respectively. Common genes were broadly associated with developmental processes **(Figure 8A & Supplementary Table S41)**; BL1 subtype was enriched mainly for hemostasis and coagulation processes^34–36;^ BL2 subtype for chemotaxis and signaling processes^37,38^; LAR subtype for metabolic process^39,40^ and lastly M subtype was associated with modification and nuclear division processes^41–43^ **(Figure 8B-E)**. Detailed discussion of the processes associated with TNBC subtype-specific mediators of resistance, and their functional relevance is discussed in **Supplementary Note 1.** Complete lists of the enriched common and unique processes in each subtype are provided in the **Supplementary Table S42-S45.** Likewise, in the case of Repressed TopNets, 6 genes -- *STAT3, TGFB1, TNF, EP300, TP53* and *FOXA1* were common in all the subtypes and were mainly associated with the transcription associated processes. 61 genes were unique in the BL1 subtype and were associated with the immune system and function. BL2 with 45 unique genes were enriched for lipid related processes; LAR with 40 unique genes were associated with signaling processes and lastly M subtype with 55 unique genes were enriched with cell division processes. Complete lists of the enriched common and unique processes in each subtype for Repressed TopNets are provided in the **Supplementary Table S46-S50 & Supplementary Figure S9A-E.** Overall, these results reveal genes potentially mediating resistance across all TNBC subtypes and also distinct processes mediating resistance in each subtype.

**Figure 8:**
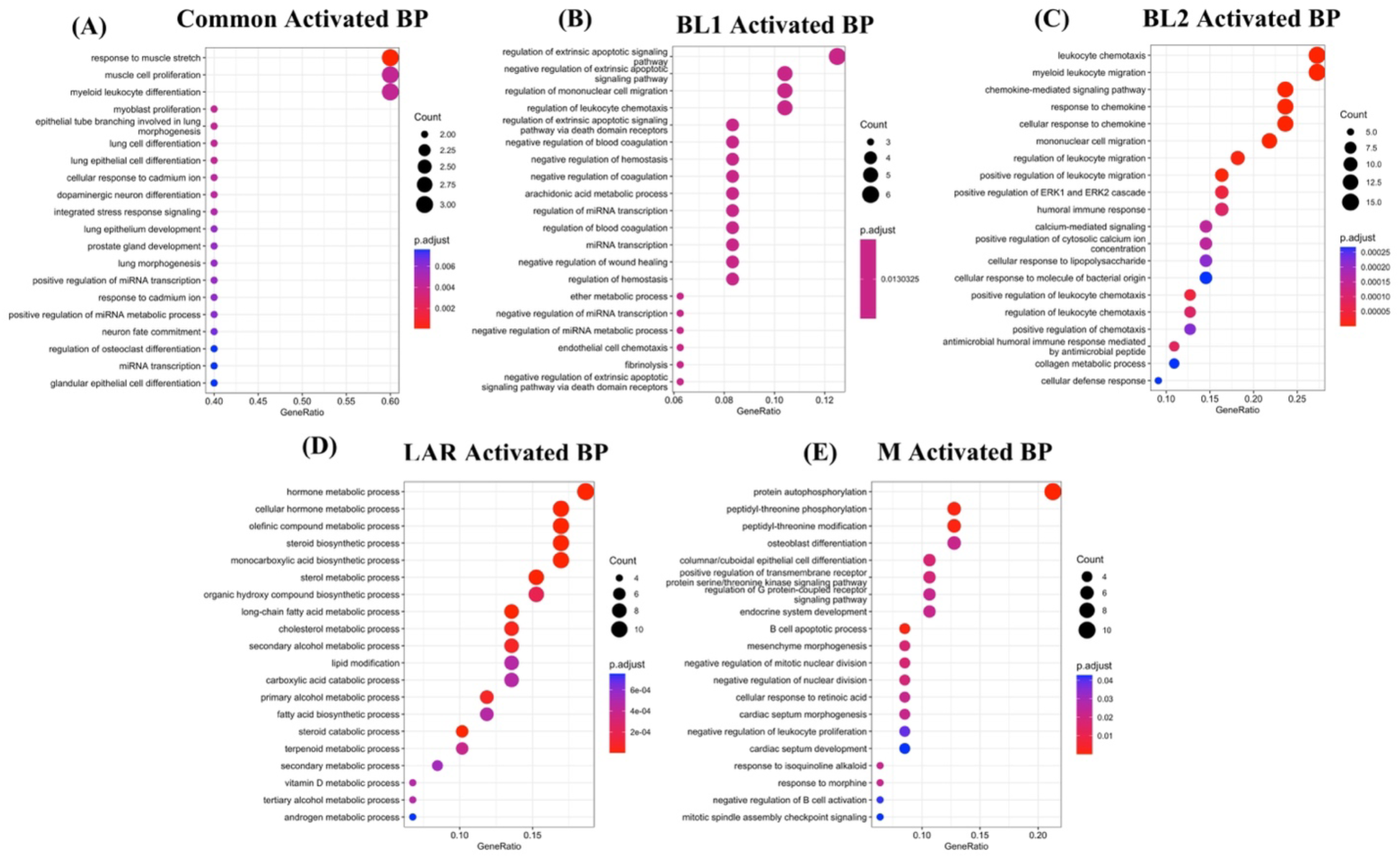
Top central PathExt Activated central genes are associated with essential biological processes in TNBC non-responders. Top20 enriched biological processes associated with common central genes among all the 4 TNBC subtypes (A). Top20 enriched biological processes associated with unique BL1 (B); BL2 (C); LAR (D); and M (E) subtype central genes.

Next, we assessed the extent to which PathExt-identified genes can help differentiate responders from non-responders. For this, we identified 21 genes common among the top 200 Activated genes in all four TNBC subtypes; for comparison, we analogously selected 13 upregulated DEGs (**Supplementary Table S51)**.

We plotted these genes’ expression to see if they can discriminate responders and non-responders in an independent dataset by Stickeler et.al (GSE21974)^44^. As shown in **Figure 9A**, PathExt-identified genes’ expression discriminate responders and non-responders with significant p-value of 0.028, however, DEGs failed to discriminate between responders and non-responders (p-value = 0.61) **(Figure 9B)**. These results underscore the generalizability of PathExt-identified genes across cohorts.

**Figure 9:**
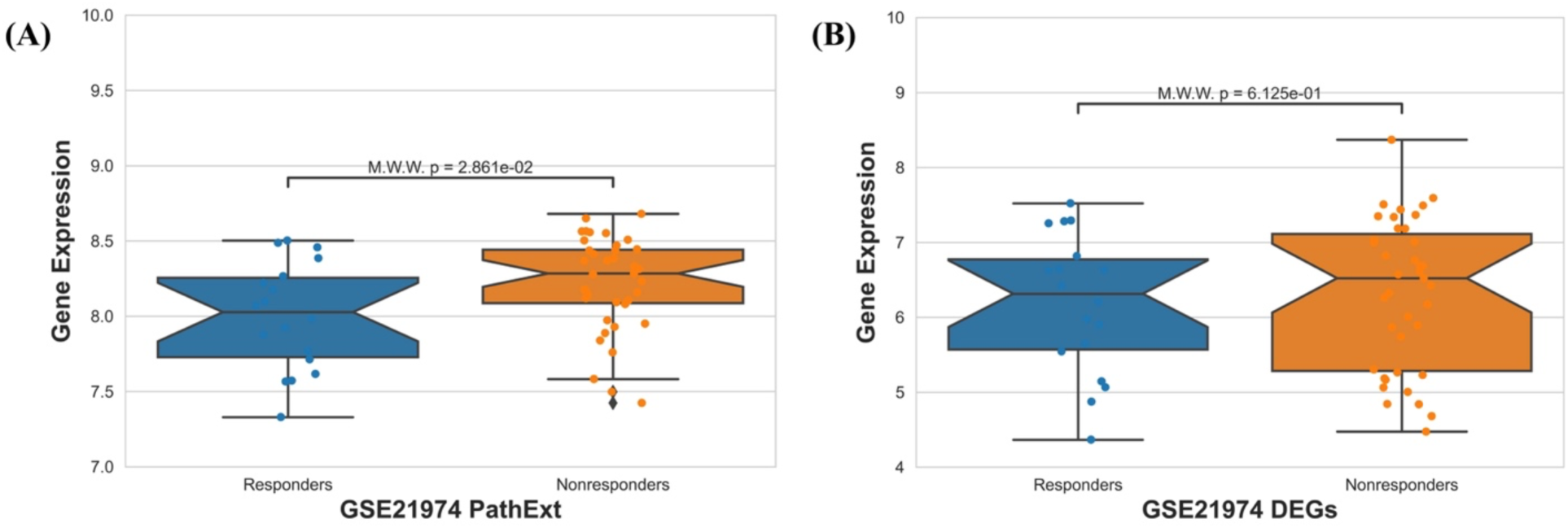
PathExt central genes discriminate responder-nonresponder significantly compared to DEGs. Average gene expression of 21 central genes from PathExt Activated TopNets and 13 upregulated DEGs were computed in each sample in an independent dataset. Boxplot show that PathExt discriminate responder and nonresponder significantly (Mann Whitney Test) and DEGs do not.

As a community resource, we perform drug repurposing analysis and provide potential drugs targeting the genes potentially mediating resistance in TNBC as currently only poly (ADP-ribose) polymerase (PARP) inhibitors and immune checkpoint inhibitors have been approved for TNBC treatment^45^. We selected top100 most frequent central genes present in the Activating TopNets of TNBC subtypes and selected the 28 genes present in at least 3 TNBC subtypes, of which 3 were excluded due to unsuitability for docking due to their structural properties. Therefore, we proceeded with 25 genes for virtual drug screening. Based on the drug repurposing tool CLUE^46^ (https://clue.io/repurposing-app), 16 of the 25 genes are targeted by at least one approved drug. **Table 1** shows the targets for which at least one drug is mapped in the CLUE data (we have provided up to 3 approved drugs mapped to each target). For the remaining 9 targets, we performed virtual screening using FDA approved drugs **(Methods; Supplementary Table S52)** and provided top 3 mapped drugs for each gene in **Supplementary Table S53**. **Supplementary Figure S10** provides an example of docking results for one of the target genes.

**Table 1:** List of the targets from the Activated TopNets and the drugs mapped in CLUE database. This table comprises of those targets for which at least one drug was mapped in CLUE database. Maximum of 3 drugs are provided for each target gene.

| Target | Top 3 potential drug |
| --- | --- |
| ALB | Erythromycin-estolate, Erythromycin-ethylsuccinate, Iodipamide |
| CALM1 | Chlorpromazine, Trifluoperazine, Loperamide |
| EGFR | Brigatinib, Olmutinib, Gefitinib |
| ESR1 | Hexestrol, Danazol, Gestrinone |
| JUN | Ephedrine-(racemic), Irbesartan, Vinblastine |
| ADCY2 | Forskolin |
| APP | Curcumin |
| CCND1 | Arsenic-trioxide |
| CCR5 | Maraviroc |
| CREB1 | Adenosine-phosphate, Naloxone |
| CTNNB1 | Urea |
| EP300 | Curcumin |
| FOS | Ephedrine-(racemic) |
| IL6 | Ibudilast |
| STAT3 | Acitretin, Niclosamide |

## Discussion

Tumor heterogeneity in breast cancer, particularly TNBC, remains an ongoing hurdle for identifying therapeutic targets with broad applicability. By identifying central mediators of gene expression changes, PathExt is designed to refine expression variance and can be used in multiple translational contexts.

PathExt achieves its goal by first identifying critical pathways in a pre-defined knowledge-based gene network, i.e., those that exhibit significantly different activity in a specific transcriptomic sample compared to controls, then identifying key mediators of the critical pathways. In doing so PathExt relies less on differential expression of individual genes (which are highly variable^47^) and more on network relationships. Unlike differential expression, which relies on sufficient sampling, PathExt can be applied to a single sample, making it applicable for clinical scenarios where large cohorts are rarer. By identifying key genes in a sample-wise fashion (as opposed to a cohort for differential expression), PathExt directly accounts for inter-sample heterogeneity. By focusing on genes mediating the global transcriptome, PathExt shows far greater commonality across BRCA subtypes across multiple benchmark sets compared to DEGs. PathExt also performs favorably to a previous network-based approach - MOMA.

We note that the central genes mediating the Activated TopNet (comprising paths with significantly higher activity in the condition of interest relative to control) are not necessarily differentially upregulated **(Supplementary Figure S11)**. Likewise, central genes mediating the Repressed TopNet (comprising paths with significantly lower activity in the condition) are not necessarily differentially downregulated **(Supplementary Figure S12)**; in particular, many of these genes may indeed have repressive effects on other genes while themselves being upregulated. This explains the somewhat counterintuitive observation that the key genes in both the Activated and the Repressed TopNets exhibit elevated inactivating mutations in cancer **(Figure 4B)**. Furthermore, the cell types expressing the genes identified based on bulk sequencing is not immediately clear. Our single-cell analysis of PathExt-identified genes points to the role of tumor microenvironment in oncogenesis, where not only the malignant but various immune compartments may play a key role in mediating the global gene expression, as profiled by bulk sequencing.

As shown above PathExt recapitulates the genes previously associated with BRCA more frequently compared to DEGs. For instance, *ERBB2* gene (which encodes for HER2) was identified as a top central gene in over 50% (42/80) of the patients by PathExt compared to fewer than 15% patients (11/80) by DEG.

Besides revealing an overall greater commonality in the key genes across samples and BRCA subtypes, PathExt also reveals shared genes and pathways between specific BRCA subtypes that are consistent with known BRCA subtype-specific biology recently reviewed by Nolan et.al.^24^. For instance, Basal and Her2 like-tumors show low and medium expression of LumA signatures respectively, and we observed that top10 most frequent LumA genes (except *TTK*, *NEK2* and *BIRC5*) were among the top genes in very few Basal tumors and in nearly half of the Her2 tumors. Equally importantly, PathExt contributes to the currently limited knowledge of subtype-specific keys genes and their associated biological processes. For instance, Tan et.al.^18^ recently showed upregulation of neuronal genes preferentially in TNBC associated with neural crest and glial development. Encouragingly, PathExt identified cell-fate determination and regulation of T-cell proliferation among the top enriched processes uniquely in basal subtype (which substantially overlaps with TNBC). Likewise, Hartman et. al.^48^ have shown high expression of *HER2* leads to secretion of interleukin-6 (*IL6)* which further activates *STAT3* ultimately contributing to tumorigenesis. Uniquely in Her2 BRCA, PathExt reveals biological processes associated with IL6 response and inflammation. Consistent with reports linking aging with luminal breast cancer^49–51^, PathExt identified “aging” as the top enriched process specifically in LumA BRCA. Focusing on subtype-specific identified genes involved in subtype-specific enriched biological processes, and further filtering them based on gene co-expression network^52^ (https://coxpresdb.jp/), we identified core subtype-specific genes worthy of future follow up. These include *MYC, MAX, E2F2* and *IL2RA* in Basal; *JAK2, ERBB2*, and *MAPK9* in Her2, and *MMP2, FGFR3, F2R, SHH* and *ADCY1* in LumA subtypes, most of which have known associations with specific BRCA subtypes.

PathExt application to TNBC response revealed 5 genes -- *FOXA1, CTNNB1, JUN, FOS* and *ALB* associated with non-responsive Activated TopNets in all four TNBC subtypes. These genes were broadly enriched for development, signaling, and transcription. One of the top enriched processes we observed associated with resistance was ‘cell fate determination’ which was also reviewed by O’Reilly et.al. as one of the factors associated with chemoresistance in TNBC^15^. *FOXA1* upregulation in basal-like cell line MDA-MB-231 leads to increased drug resistance^53^. Likewise, *CTNNB1* is associated with Wnt signaling pathways, known to be associated with BRCA^54^. There is literature support for other genes as well^55,56^. For the Repressed TopNets, 6 genes were common among all the subtypes, viz. *STAT3, TGFB1, TNF, EP300, TP53* and *FOXA1.* Interestingly, *FOXA1* is revealed as a key gene in both Activated and Repressed TopNets across all subtypes. Dai et.al. have shown that downregulation of *FOXA1* leads to increased malignancy and cancer stemness by suppressing *SOD2* and *IL6*^57^, underscoring a pleiotropic role of *FOXA1*. Another resistance-associated gene PathExt revealed is *EP300*, a known modulator of paclitaxel resistance and stemness^58^. Interestingly, the study we used for our analysis included Paclitaxel treatment after neoadjuvant chemotherapy.

Overall, PathExt is complementary to the conventional DEG approach and reveals common and BRCA subtype-specific key genes and processes, as well as gene mediating chemotherapy response in TNBC. Lastly, as a community resource, we have provided potential drugs that are either approved or undergoing trials targeting novel key genes revealed by our analyses.

## Methods

### Data Collection and processing

Transcriptomic profiles for 1059 primary BRCA tumors and 112 normal adjacent samples were downloaded from TCGA^59^. Gene expression was jointly quantile normalized and was used as an input for the PathExt^60^. We also analyzed the TNBC transcriptomic dataset by Horak et.al.^33^ where they look for the biomarkers associated with the response to the neoadjuvant doxorubicin/cyclophosphamide followed by Ixabepilone. In a follow up study^61^, Lehmann et.al. classified 138 of the above TNBC samples into 4 TNBC subtypes (see ^61^ for classification details) - BL1 (47 samples), BL2 (27 samples), LAR (21 samples), and M (43 samples). We applied PathExt to each TNBC subtype separately to identify central genes mediating resistance (non-response).

### Identification of sample-specific central genes using PathExt

PathExt method has been described in detail in our previous publications^10,60^. Here we provide a very brief sketch. Based on a set of control transcription profiles (e.g., normal adjacent breast samples for a BRCA sample), we first compute weight for each node in a given knowledge-based protein interaction network, such that the node weight represents the extent of upregulation (Activated) or downregulation (Repressed) of the gene in the foreground sample relative to the control. PathExt then identifies significantly perturbed paths in the network and finally in the subnetwork composed of such significantly perturbed paths (Activated or Repressed TopNet), it then identifies the top central genes based on betweenness centrality measure^62^. These central genes represent key mediators of global transcription changes (upregulation for Activated TopNet and downregulation for Repressed TopNet) in a particular sample. Having computed the top 100 genes for each BRCA sample we compute the number of sample (of a given BRCA subtype) in which a gene was among the central genes, and then selected the top 200 most frequent genes to represent subtype-specific key genes; this was done independently for Activated and Repressed TopNets. For comparison, based on log-fold change for each gene, we selected top 100 up and downregulated genes for each sample and further based on frequency we selected top 200 genes (up and downregulated) from each subtype. Analogous procedure was applied to identify key genes corresponding to non-responders relative to responders among the TNBC BRCA cohort.

### Gene Ontology Enrichment Analysis

To identify enriched biological processes and molecular functions in various subtypes, we used the Clusterprofiler 4.0 package^63^, where the foreground was the top200 central genes (PathExt or DEGs) and the background was the default background used by the package. The database used was the human database and the minimum gene size and maximum gene size was set as 10 and 200 respectively to ensure specific terms. We then used the function “simplify” with cutoff value 0.8 to remove the redundant terms. Default parameters were used otherwise. In the case of TNBC responder and non-responders, we used similar parameters except foreground and background gene list. In this case, the foreground was top100 genes and the background was a customized list of 10994 genes, those that were detected in the cohort **[Supplementary Table S54]**

### Cell-line specific genetic dependency

From the DepMap database^13^, we obtained the gene dependency probability scores for 16708 genes in 44 breast cancer cell lines, further categorized into BRCA subtypes. For each gene, we then obtained the average dependency probability score across the cell lines in a subtype-specific manner. As recommended, genes with average dependency probability score >= 0.5 were recorded as essential for comparing PathExt and DEG genes.

### Comparison of PathExt and DEGs genes with benchmarks gene sets

We compiled various previously published datasets of potential BRCA driver genes, approved FDA drugs for BRCA and assessed their overlap with the PathExt central genes or DEGs. We used Fisher exact test to assess the statistical significance of overlap and reported Odds Ratio (>1 indicates greater than expected overlap).

### Breast cancer subtype single cell data analysis

We used a single cell RNA-seq dataset for the 4 BRCA subtypes by Qian et.al.^31^. Read count matrices of scRNA-seq data (obtained using 10X v2 sequencing) breast cancer tumors were downloaded from http://blueprint.lambrechtslab.org. Author-supplied annotations were used to label each cell. The miQC package^64^ was used to remove dead cells, with a probability threshold of 0.5 (i.e., a probability of at least 0.5 that the cell is not dead) used to retain high-quality cells for downstream analyses. After filtering the dataset, we retained only those cell types having a minimum of 50 cells. This provided 7 cell type data for Basal and Her2 subtypes, and 4 cell type data for LumA & LumB subtypes. Next, we computed the z-score of gene expression for the PathExt top200 central genes in each cell (using all cells to estimate the mean and the standard deviation) and then took the average z-score value of the cells in a given cell type. Next, we selected the genes with z-score > 0 in each cell type and were considered as ‘expressed’ in that cell type. Next, we computed the fraction (%) of such expressed genes in every cell type and normalized the fraction across all cell types to get ‘Observed frequency’ in a cell type. We also computed the ‘Expected frequency’ in a similar manner by considering all genes. Finally, Log(Observed/Expected) value was computed for the central genes across cell types.

### Identifying key genes associated with neoadjuvant treatment in TNBC subtypes

PathExt was applied to a previously published cohort of TNBC tumors having undergone neoadjuvant therapy with Doxorubicin/Cyclophosphamide followed by Ixabepilone or Paclitaxel drug, and further classified into responders and non-responders^33^ where responders include patients with complete or partial response, whereas non-responders include patients with stable or progressive disease. TNBC is further subdivided into 4 subtypes, BL1, BL2, LAR and M^61^, and hence, we applied PathExt in a subtype-specific manner to identify central genes among non-responders relative to responders.

### Drug repurposing study

We selected the top 100 most frequent central genes present in Activated TopNets from each TNBC subtype and then further selected genes present in at least 3 subtypes. This resulted in 29 genes out of which 3 genes were not suitable for analysis because of their structural properties. Based on the CMap database (using CLUE platform)^65^, we first identified the drugs targeting these genes. We consider only those drugs which are either approved or undergoing trials. For the remaining unmapped targets, we performed virtual screening using drugs which are either FDA approved or are under clinical trials. The 3D structures of the targets were downloaded from the RCSB-PDB^66^ and further refined using OpenBabel software^67^. Next, we used online server DrugRep^68^ to locate various binding pockets on the protein surface and selected the pocket with the largest volume covering most active site residues. Identification of active sites and residues facilitate the docking experiment and help in identifying drugs with improved binding. Virtual Screening was performed using Autodock Vina software^69^. The software requires a receptor and ligand file in a specific file format (“pdbqt” file format), which was prepared using MGLTools software^70^. Post virtual screening, based on docking free energy and root mean square deviation (RMSD), top 3 drugs for each target were proposed. Further, experimental work is required to ^66^prove the efficacy of these drugs.

## Acknowledgement

This work utilized the computational resources of the NIH HPC Biowulf cluster.

## Author contribution

PA and NJ download and processed the data. VG downloaded and processed the single cell data. PA, AT, AS and SH perform the analysis. PA and SH perform the statistical analysis. PA, PSR and SH wrote the manuscript. PA and SH supervised the study. All authors read the article and approved the submitted version.

## Data Availability Statement

The original contributions presented in the study are included in the article/Supplementary Material. Further inquiries can be directed to the corresponding authors.

