## Supplementary Figures for "Network-based approach elucidates critical genes in BRCA subtypes and chemotherapy response in Triple Negative Breast Cancer"

* Equal Correspondence

**Supplementary Figures**

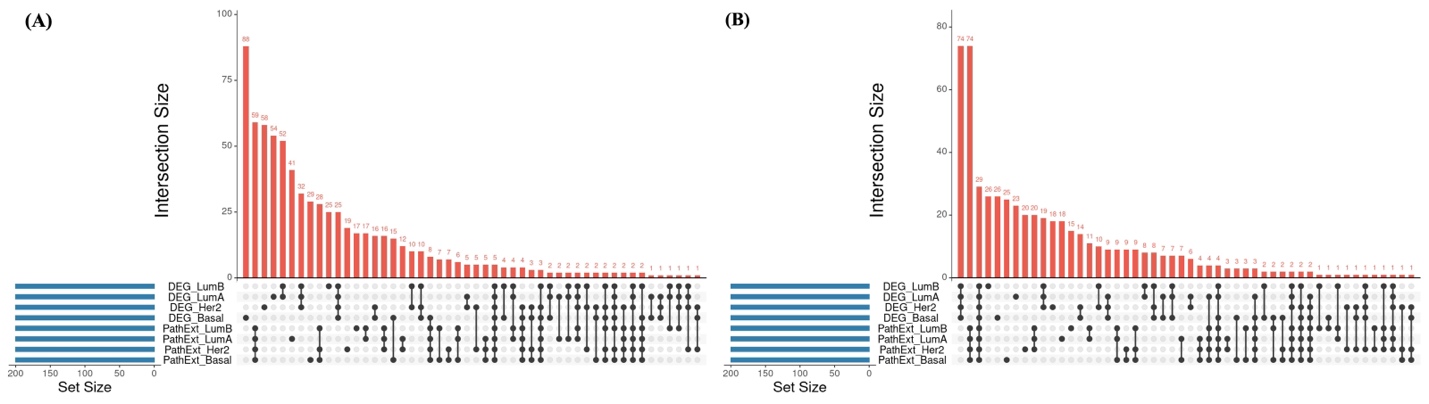

**Figure S1: PathExt and DEGs do not share similar central genes.** Upset Plot showing commonality among PathExt and DEG subtypes for (A) Activated TopNets and Upregulated genes and (B) Repressed TopNet and downregulated genes.

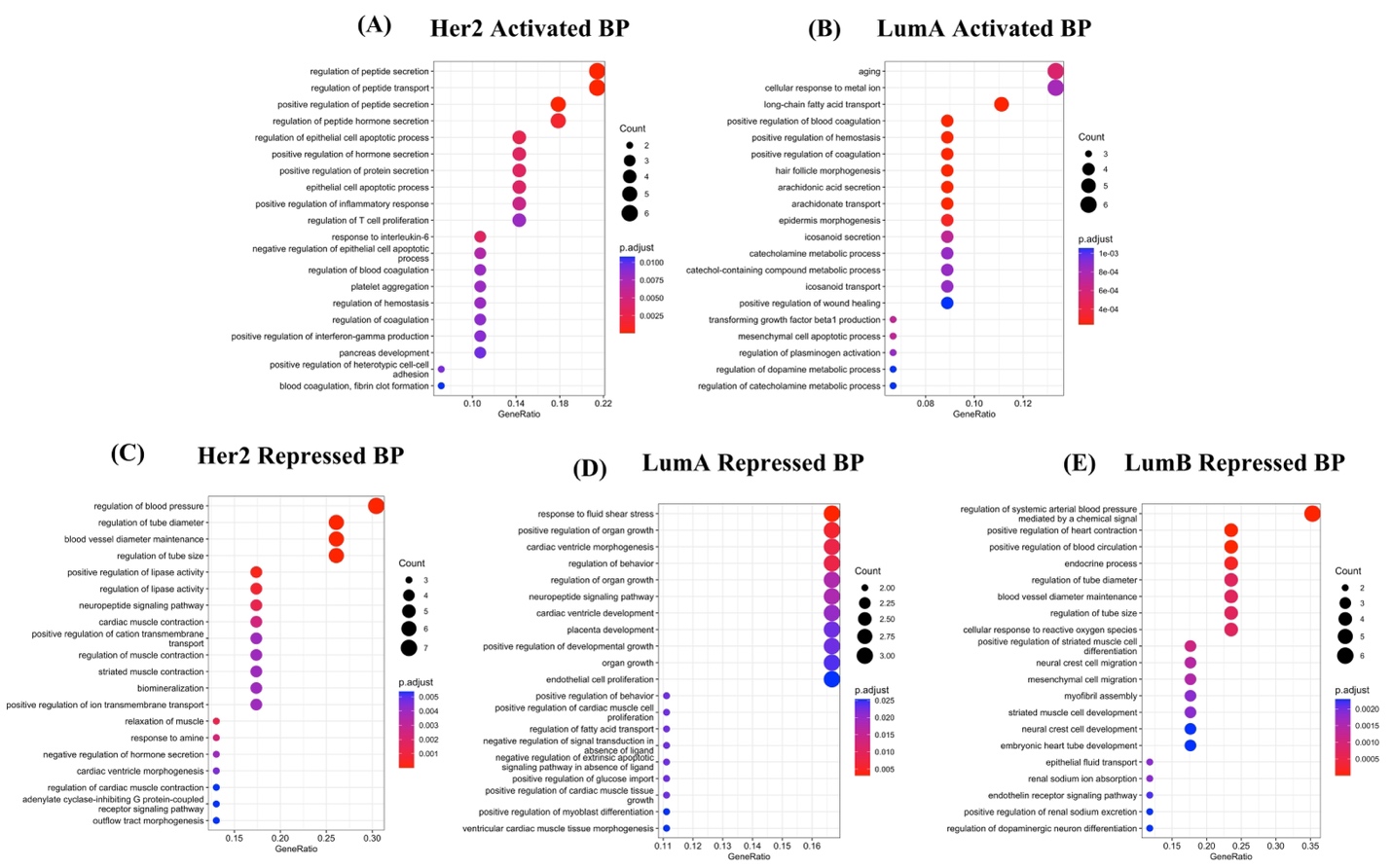

**Figure S2: Top central PathExt central genes are associated with essential biological processes.** Top20 enriched biological processes associated with activated genes uniquely in Her2 (A); Top20 enriched biological processes associated with activated genes uniquely in Luminal-A (B); Top20 enriched biological processes associated with repressed genes uniquely in Her2 (C); Top20 enriched biological processes associated with repressed genes uniquely in Luminal-A (D); and Top20 enriched biological processes associated with repressed genes uniquely in Luminal-B (E).

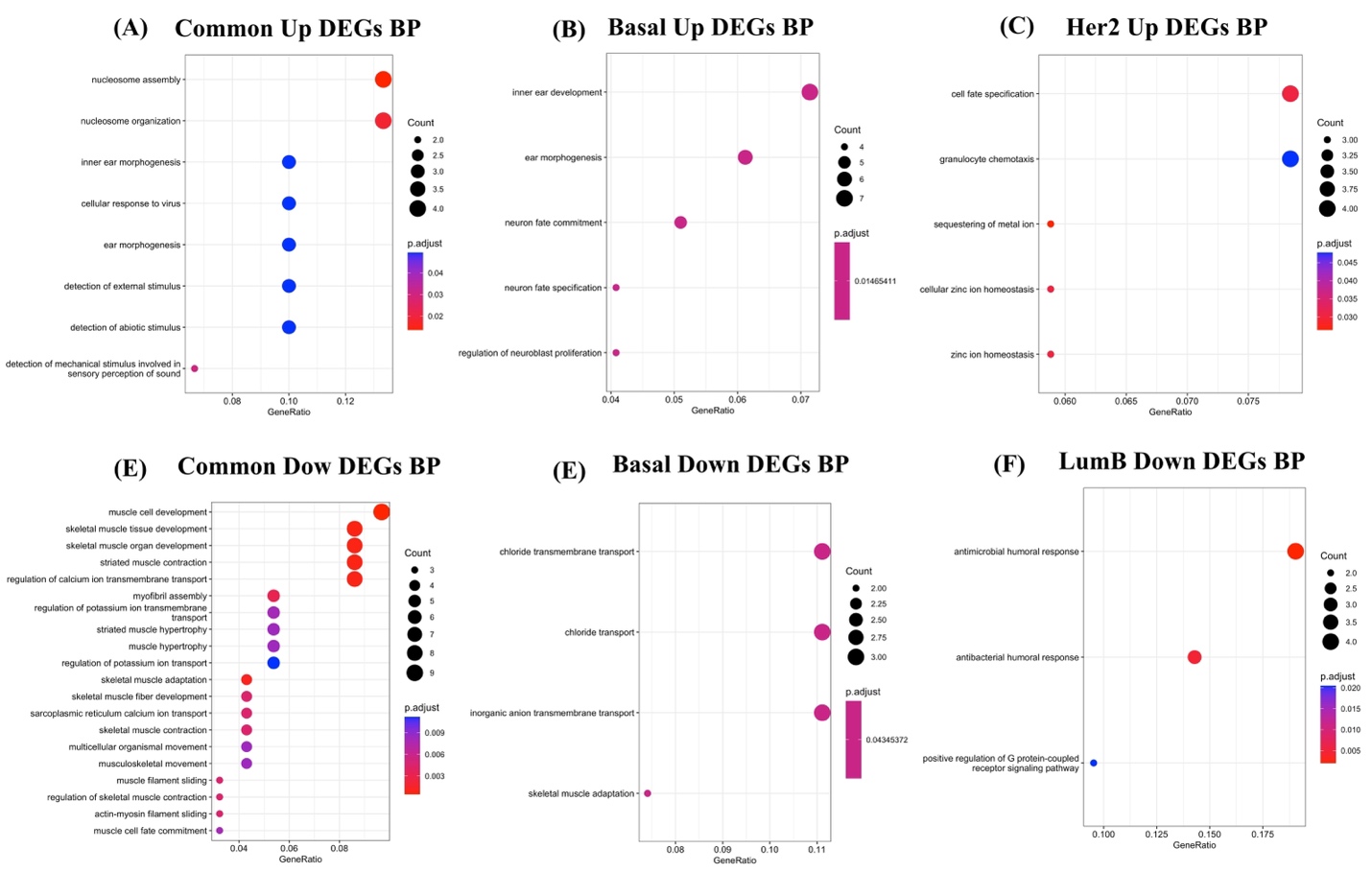

**Figure S3: Top central DEGs are associated with few biological processes.** Enriched biological processes associated with common upregulated central DEGs among all the 4 BRCA subtypes (A); unique upregulated central DEGs in Basal subtype (B); upregulated central DEGs in Her2 subtype (C); common downregulated central DEGs among all the 4 BRCA subtypes (D); unique downregulated central DEGs in Basal subtype (E), and unique downregulated central DEGs in LumB subtype (F).

*** ‘BP’ in the figure denotes Biological Processes**

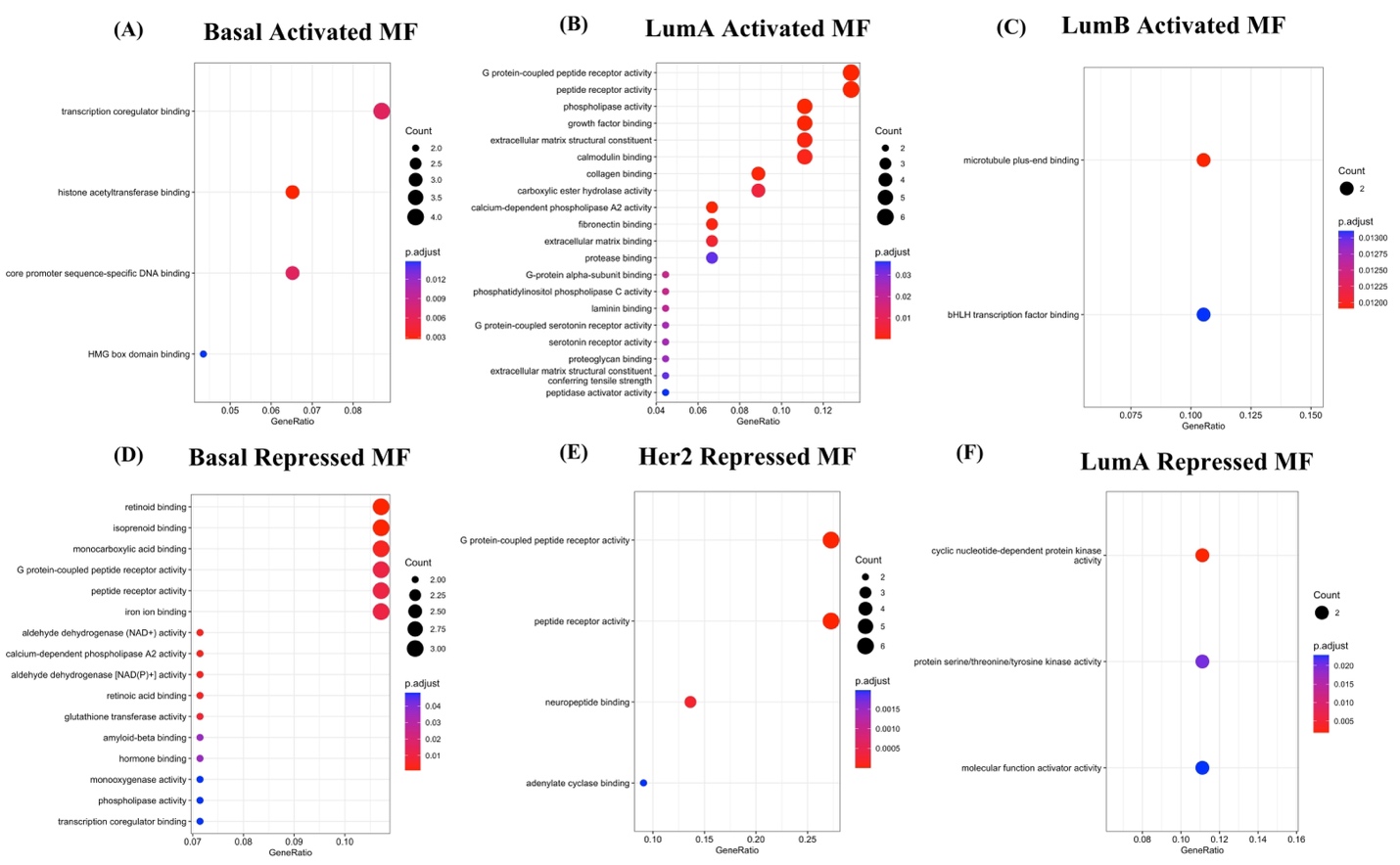

**Figure S4: Top central PathExt genes (Activated & Repressed) are associated with essential molecular functions.** Enriched molecular functions associated with Activated Basal unique genes (A); Activated Luminal-A unique genes (B); Activated Luminal-B unique genes (C); Repressed Basal unique genes (D); Repressed Her2 unique genes (E); and Repressed Luminal-A unique genes (F).

*** ‘MF’ in the figure denotes Molecular Functions**

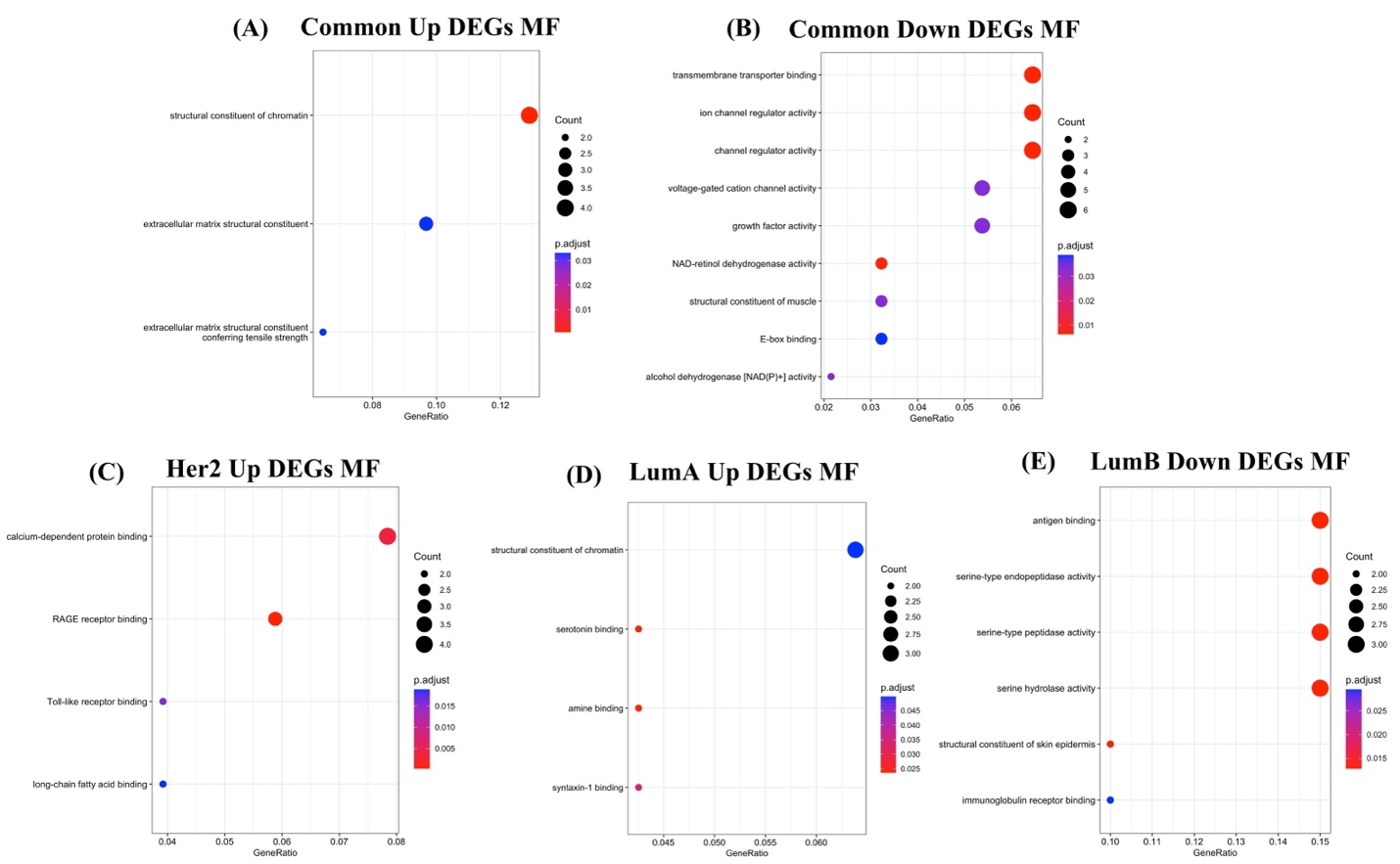

**Figure S5: Top central PathExt genes (Activated & Repressed) are associated with essential molecular functions.** Enriched molecular functions associated with Pan-common subtype Upregulated DEGs (A); Pan-common subtype Downregulated DEGs (B); Upregulated Her2 Unique central genes (C); Upregulated LumA Unique central genes (D); and Downregulated LumB Unique central genes (E).

*** ‘MF’ in the figure denotes Molecular Functions**

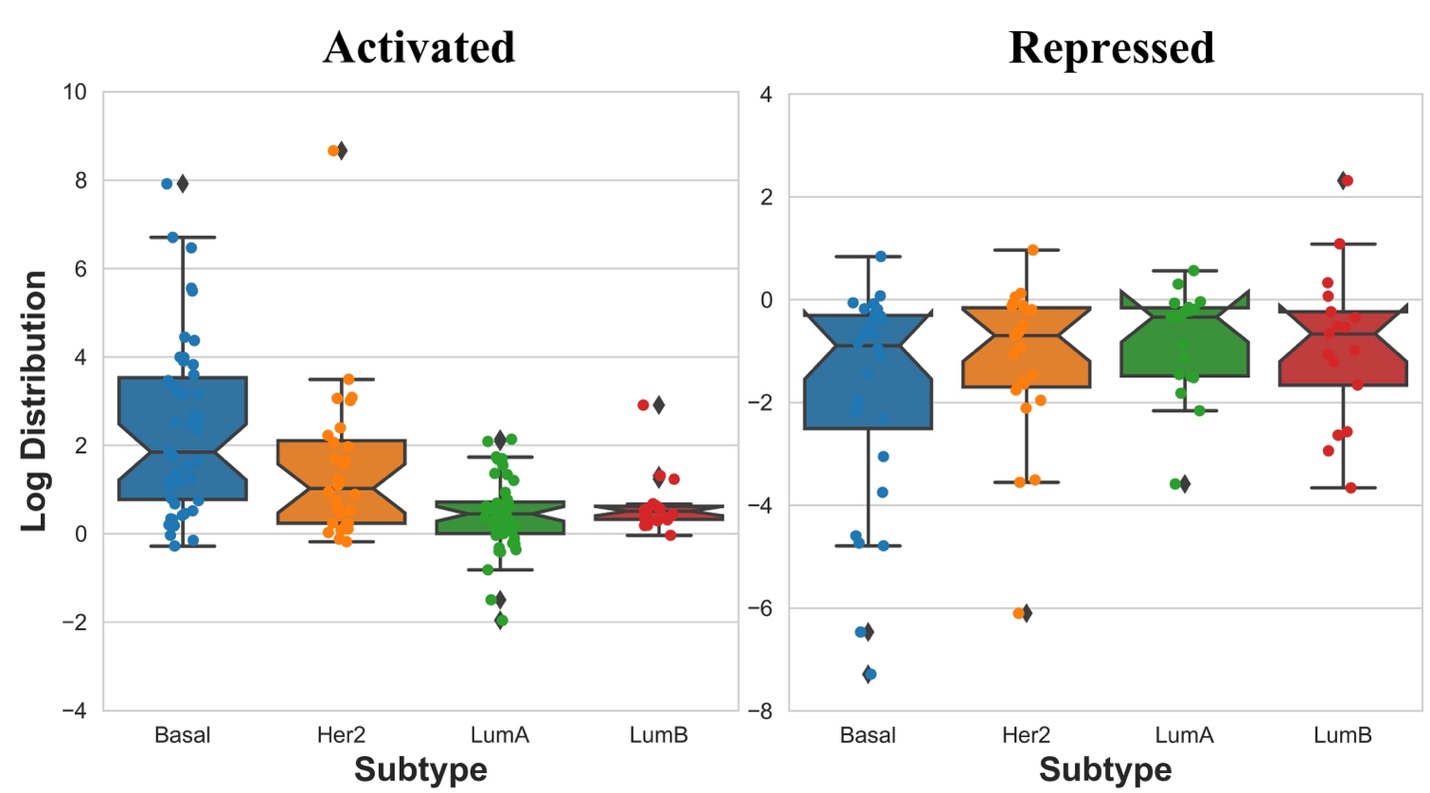

**Figure S6: PathExt reveals subtype specific gene expression properties.** Boxplot representation of subtype specific PathExt Activated and Repressed TopNets unique gene expression

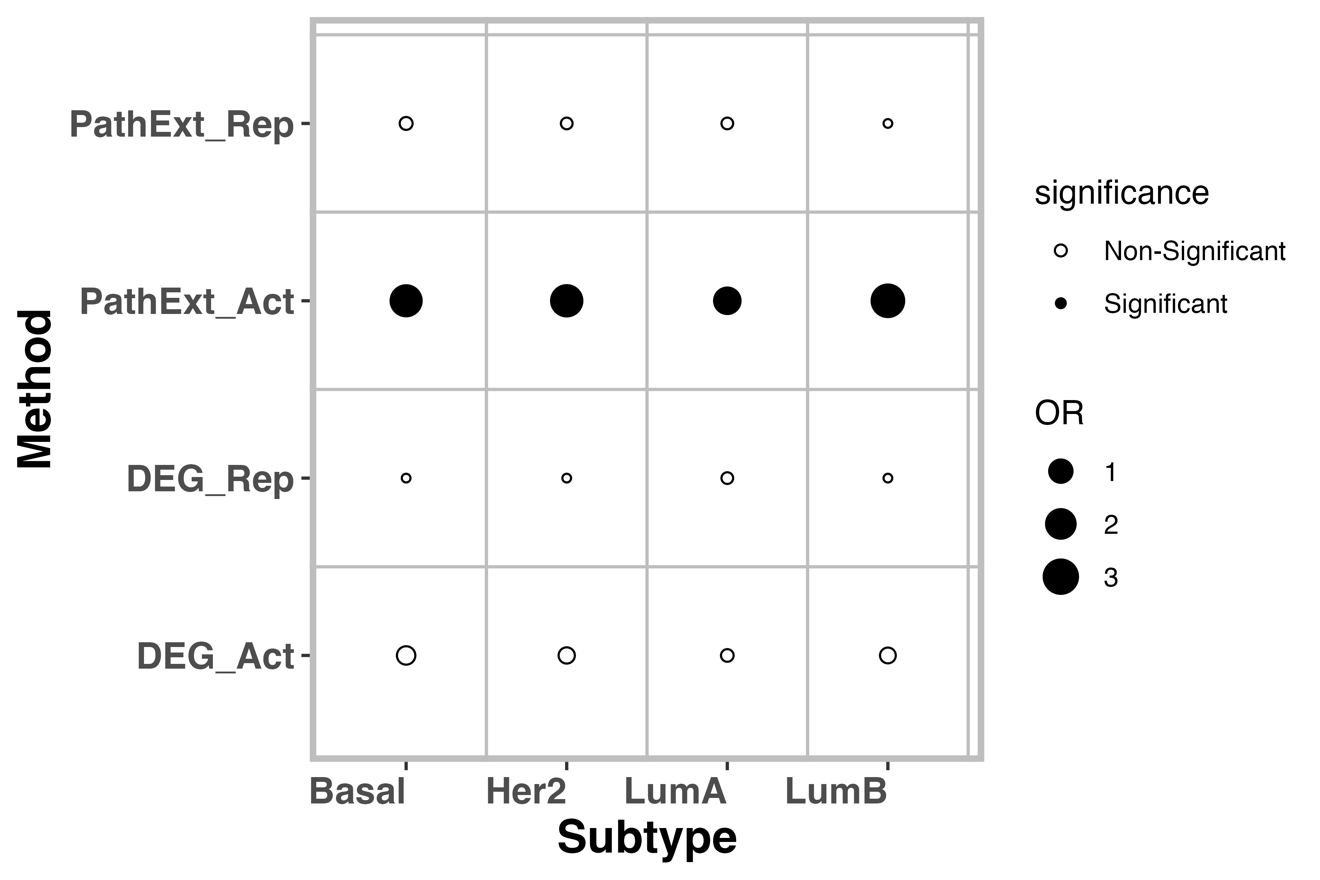

**Supplementary Fig S7:** Dotplot representation of Fisher’s Odds Ratio showing the significance of recapitulation of genes with dependency probability >=0.5 in by PathExt and DEGs central genes.

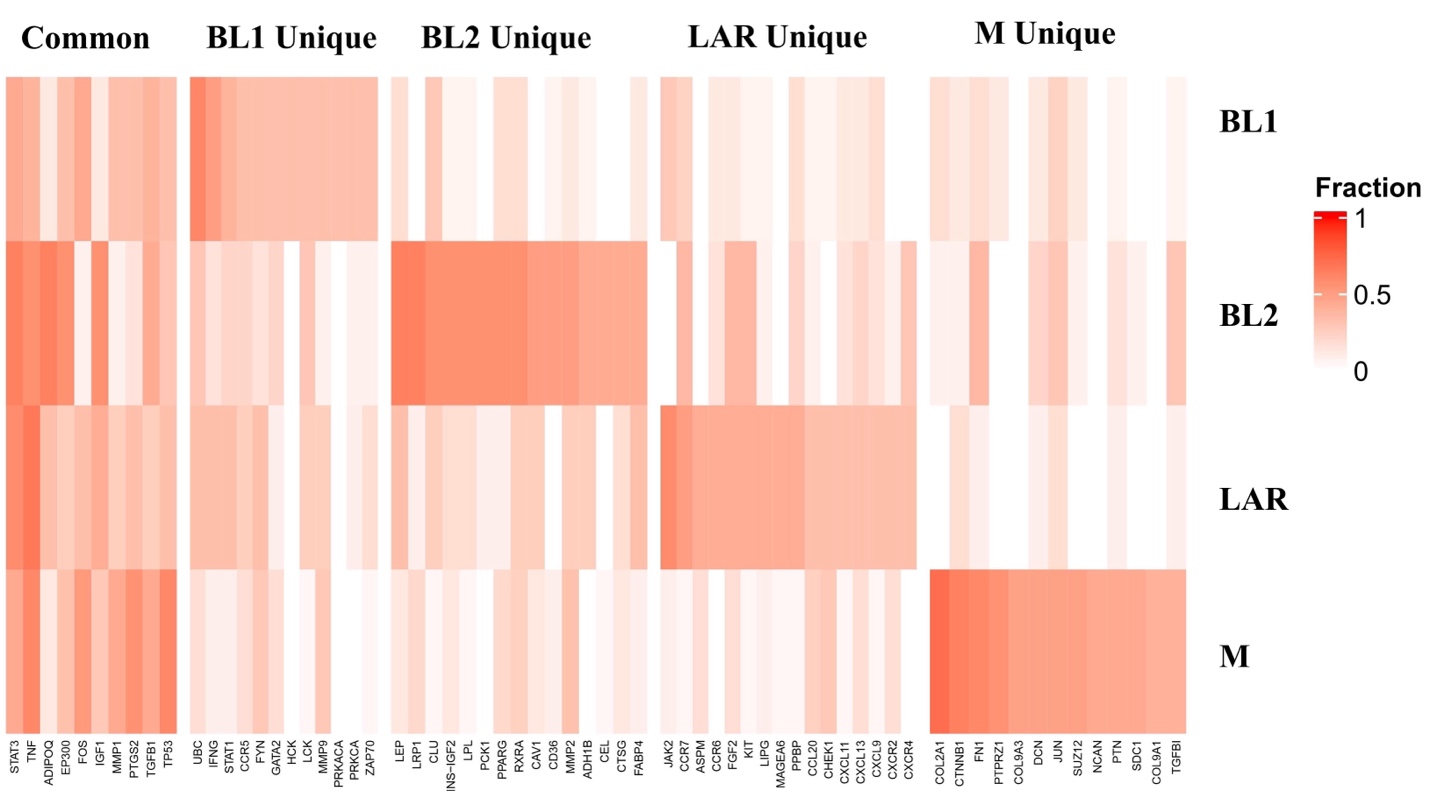

**Figure S8:** **Top PathExt Repressed unique and common gene fraction distribution in various TNBC subtypes**. Top20 most frequent genes were selected from each TNBC subtype, from which pan-subtype common genes (top20 in at least two subtypes) and subtype-specific unique genes were identified. The heat plot shows, in each subtype, the fraction of samples in which the gene was among the top 20 central genes.

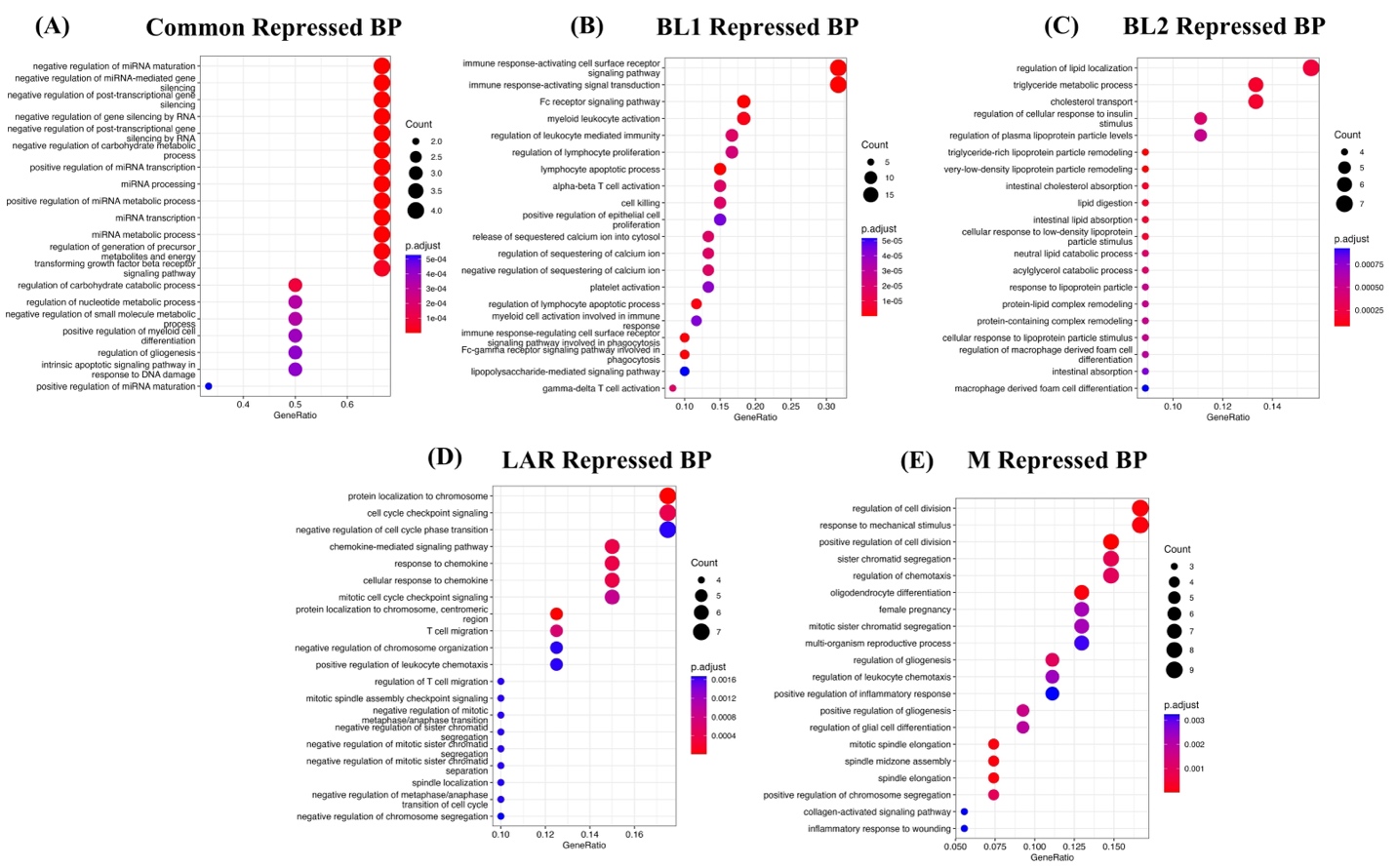

**Figure S9: Top central PathExt Repressed central genes are associated with essential biological processes in TNBC non-responders.** Top20 enriched biological processes associated with common central genes among all the 4 TNBC subtypes (A). Top20 enriched biological processes associated with unique BL1 (B); BL2 (C); LAR (D); and M (E) subtype central genes.

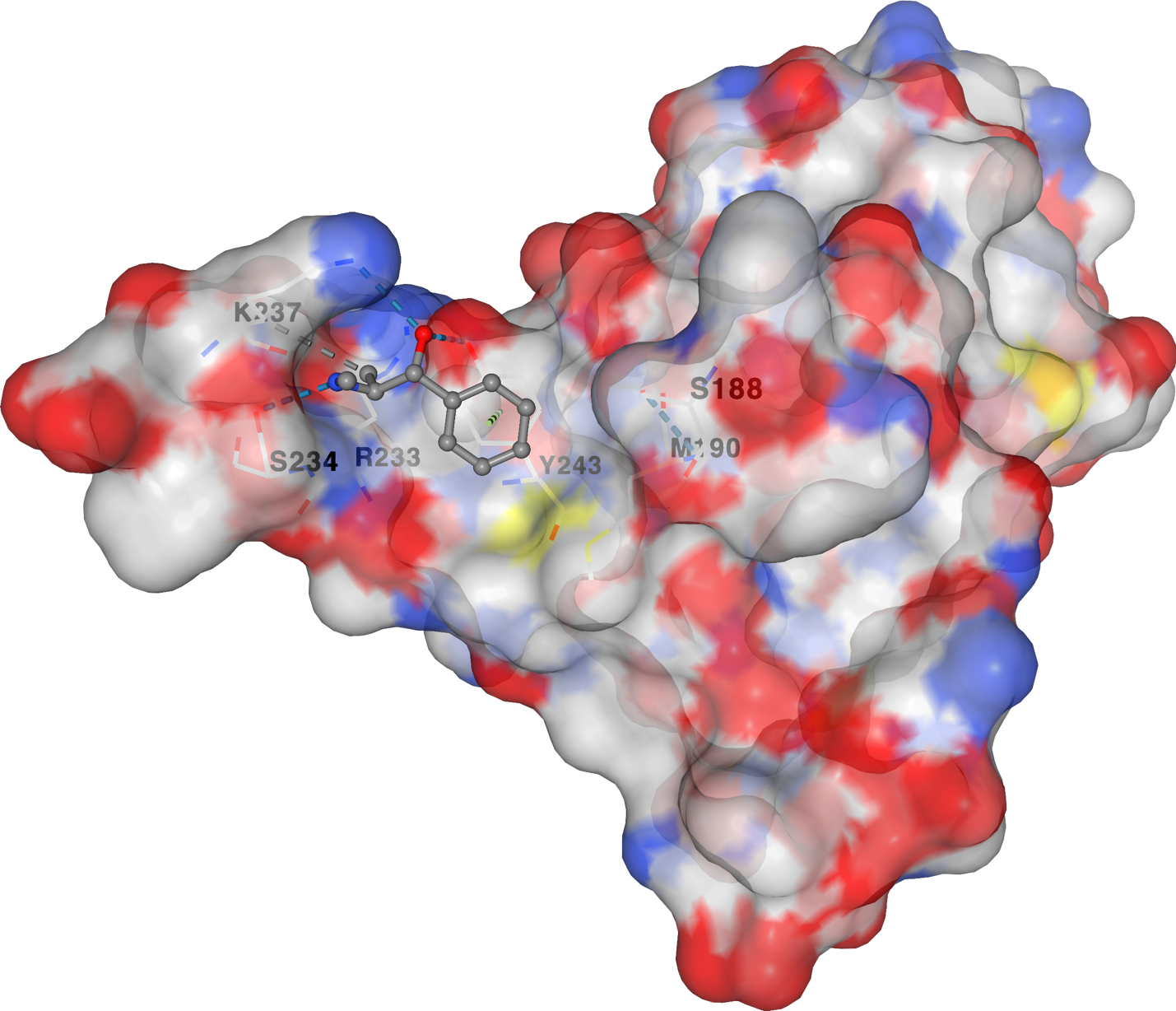

**Figure S10:** **Docking complex of FOXA1 protein with drug Ephedrine in the active site region.** We performed docking experiments using CBDock2 online server (<https://cadd.labshare.cn/cb-dock2/php/blinddock.php>) and we can see the strong interaction of the target residues (S188, M190, R233, S234, K237 and Y243) with the drug.

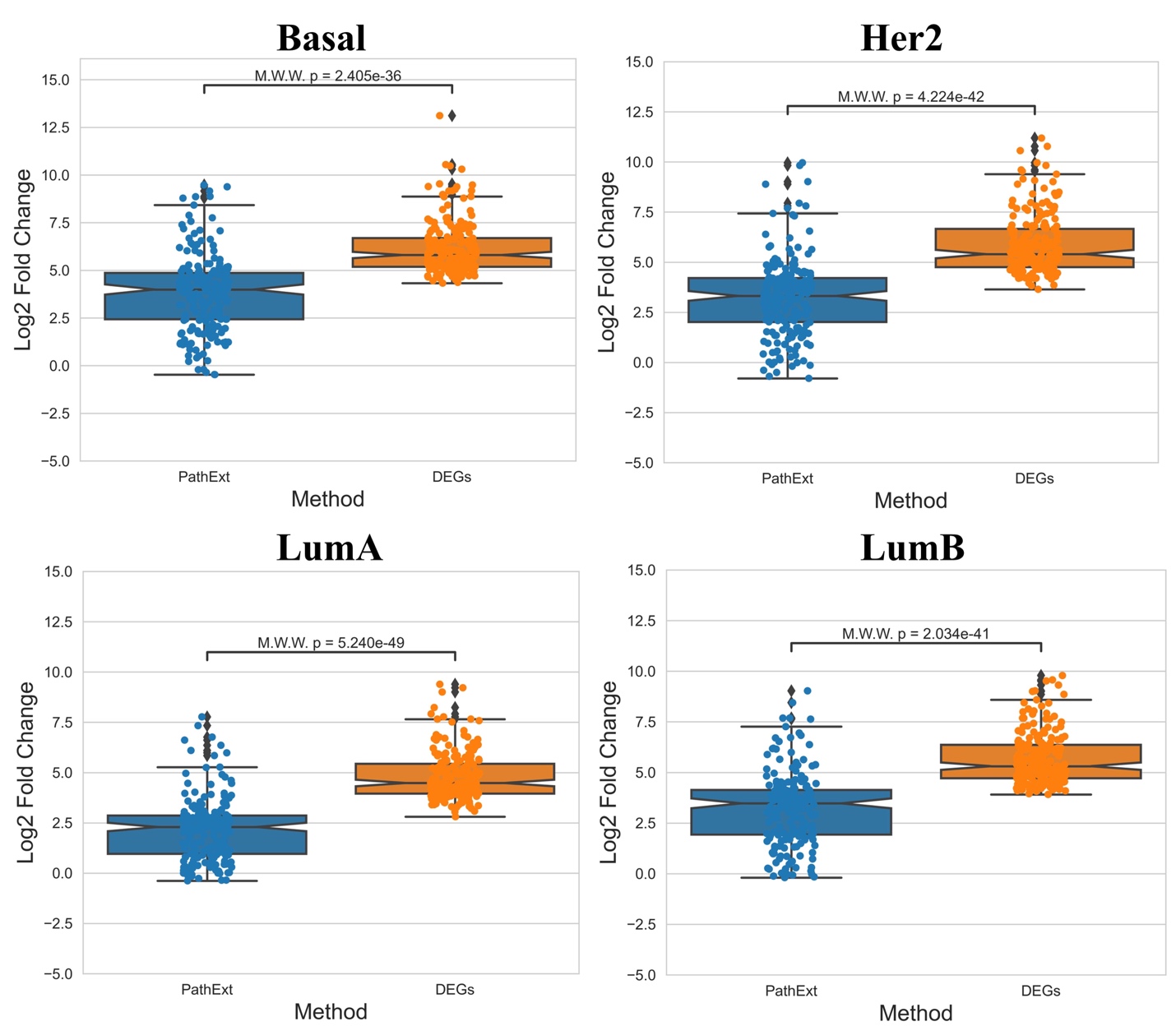

**Figure S11: Gene Expression LogFC comparison between PathExt Activated Central genes and DEGs upregulated central genes.** Log Fold change using mean gene expression of the top200 central genes and DEGs was computed showing the lower fold change of the PathExt genes compared to upregulated DEGs for Basal (A); Her2 (B); Luminal-A (C); and Luminal-B (D) subtype.

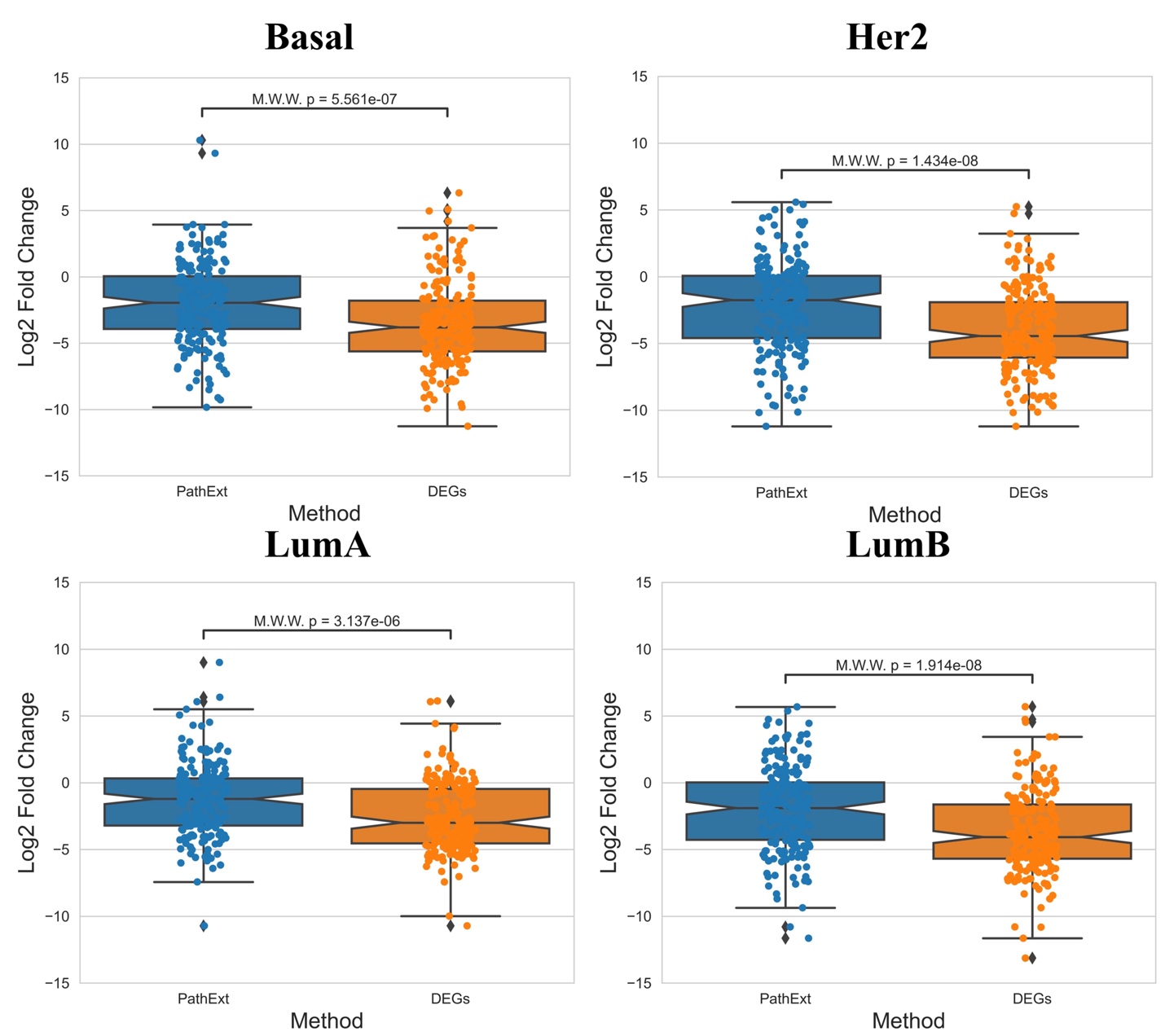

**Figure S12: Gene Expression LogFC comparison between PathExt Repressed Central genes and DEGs downregulated central genes.** Log Fold change using mean gene expression of the top200 central genes and DEGs was computed showing the lower fold change of the PathExt genes compared to downregulated DEGs for Basal (A); Her2 (B); Luminal-A (C); and Luminal-B (D) subtype.

**Supplementary Note 1: Processes enriched among central genes in Activated TopNet in TNBC Non-responders.**

We examine the most frequent top 100 genes in each subtype, for the Activated TopNets, 5 genes -- *FOXA1, CTNNB1, JUN, FOS* and *ALB* were found in all TNBC subtypes. These genes are associated prominently with differentiation, stress response and *TGFβ* signaling, etc. **(Fig 8A & Supplementary Table S40)**. Multiple studies have shown that *FOXA1* function as prognostic marker in various subtypes including TNBC where it can either co-express with Androgen Receptor (*AR*) **[PMID 29880907]** or it can transcriptionally silence *SOD2* and *IL6* **[PMID 31182923].** Likewise, *CTNNB1* has been shown to be involved in Wnt Signaling Pathway and it has been shown that level of *CTNNB1* in WNT/CTNNB1 signaling is elevated in TNBC and has poor clinical prognoses **[PMID 20395444, 33664239].** *c-JUN* activation has been associated with many cellular processes and one of the examples is the activation of *JNK* (c-Jun N-terminal kinase). And the high level of *JNK* has been observed to be elevated in TNBC **[PMID 27941886].** Likewise, role of *FOS* and *ALB* has also been reported in TNBC **[PMID 34389675, 34908845].**

Interestingly, subtype-specific genes (based on top 100) were enriched for distinct processes **(Fig 8B-E)** and were also supported by literature to a large extent. For example, BL1 was enriched for regulation of apoptotic signaling, leukocyte chemotaxis, and wound healing, all of which have been shown to associate with immune evasion and therapy resistance **[PMID 16892092, 33912188, 19527773].** Likewise, BL2 was mainly associated with humoral immune response and immune cell migration **[PMID 15686628, 31408436].** BL2 was enriched for genes encoding chemokines for example *CXCL8*, *CXCL1*, etc. These chemokines are associated with directing immune cell migration necessary to mount and delivering an effective anti-tumor immune response **[PMID 33838745]**; LAR was highly enriched for metabolic processes (hormone metabolic, Vitamin D metabolic) and lipid modification **[PMID 32296646, 32456160].** Gong et al. classified the TNBC samples into 3 categories based on their metabolic features and their further analysis reveals that inhibiting lactate dehydrogenase could enhance the response to anti-PD1 immunotherapy **[PMID 33181091].** Lastly, M subtype was associated with protein modification (protein autophosphorylation, peptidyl-threonine modification), signaling (GPCR signaling pathway, mitotic spindle checkpoint signaling), and negative regulation of nuclear division **[PMID 36077757, 25981295, 35965791].**
